# A single-nucleus and spatial transcriptomic atlas of the COVID-19 liver reveals topological, functional, and regenerative organ disruption in patients

**DOI:** 10.1101/2022.10.27.514070

**Authors:** Yered Pita-Juarez, Dimitra Karagkouni, Nikolaos Kalavros, Johannes C. Melms, Sebastian Niezen, Toni M. Delorey, Adam L Essene, Olga R. Brook, Deepti Pant, Disha Skelton-Badlani, Pourya Naderi, Pinzhu Huang, Liuliu Pan, Tyler Hether, Tallulah S. Andrews, Carly G.K. Ziegler, Jason Reeves, Andriy Myloserdnyy, Rachel Chen, Andy Nam, Stefan Phelan, Yan Liang, Amit Dipak Amin, Jana Biermann, Hanina Hibshoosh, Molly Veregge, Zachary Kramer, Christopher Jacobs, Yusuf Yalcin, Devan Phillips, Michal Slyper, Ayshwarya Subramanian, Orr Ashenberg, Zohar Bloom-Ackermann, Victoria M. Tran, James Gomez, Alexander Sturm, Shuting Zhang, Stephen J. Fleming, Sarah Warren, Joseph Beechem, Deborah Hung, Mehrtash Babadi, Robert F. Padera, Sonya A. MacParland, Gary D. Bader, Nasser Imad, Isaac H. Solomon, Eric Miller, Stefan Riedel, Caroline B.M. Porter, Alexandra-Chloé Villani, Linus T.-Y. Tsai, Winston Hide, Gyongyi Szabo, Jonathan Hecht, Orit Rozenblatt-Rosen, Alex K. Shalek, Benjamin Izar, Aviv Regev, Yury Popov, Z. Gordon Jiang, Ioannis S. Vlachos

**Author notes:** These authors contributed equally (Yered Pita-Juarez, Dimitra Karagkouni, Nikolaos Kalavros, Johannes C. Melms, Sebastian Niezen, Toni M. Delorey). Co-senior authors (Deborah Hung, Mehrtash Babadi, Robert F. Padera, Sonya A. MacParland, Gary D. Bader, Nasser Imad, Isaac H. Solomon, Eric Miller, Stefan Riedel, Caroline B.M. Porter, Alexandra-Chloé Villani, Linus T.-Y. Tsai, Winston Hide, Gyongyi Szabo, Jonathan Hecht, Orit Rozenblatt-Rosen, Alex K. Shalek, Benjamin Izar, Aviv Regev, Yury Popov, Z. Gordon Jiang, Ioannis S. Vlachos).

## Abstract

The molecular underpinnings of organ dysfunction in acute COVID-19 and its potential long-term sequelae are under intense investigation. To shed light on these in the context of liver function, we performed single-nucleus RNA-seq and spatial transcriptomic profiling of livers from 17 COVID-19 decedents. We identified hepatocytes positive for SARS-CoV-2 RNA with an expression phenotype resembling infected lung epithelial cells. Integrated analysis and comparisons with healthy controls revealed extensive changes in the cellular composition and expression states in COVID-19 liver, reflecting hepatocellular injury, ductular reaction, pathologic vascular expansion, and fibrogenesis. We also observed Kupffer cell proliferation and erythrocyte progenitors for the first time in a human liver single-cell atlas, resembling similar responses in liver injury in mice and in sepsis, respectively. Despite the absence of a clinical acute liver injury phenotype, endothelial cell composition was dramatically impacted in COVID-19, concomitantly with extensive alterations and profibrogenic activation of reactive cholangiocytes and mesenchymal cells. Our atlas provides novel insights into liver physiology and pathology in COVID-19 and forms a foundational resource for its investigation and understanding.

## Main

COVID-19 exhibits a wide phenotypic spectrum with potential multi-organ involvement during its acute phase ^1^, including liver-related pathology. Abnormal liver biochemistry is reported in 15-65% of SARS-CoV-2 infected individuals ^2–4^, and is often associated with poorer clinical outcomes ^3,4^. To date, there are few studies of human liver tissue from COVID-19 patients, hindering in-depth investigations of COVID-19-related liver injury, its main causes, and potential long-term effects, especially post-acute sequelae of SARS-CoV-2 infection (PASC), such as the patient-coined term “long COVID” (Crook et al. 2021) and post-COVID cholangiopathy, an emerging entity that may require liver transplantation ^5^. In our previous work ^6,7^, we assembled a multi-tissue COVID-19 cell atlas across lung, heart, kidney, and liver, collected at autopsy from patients who succumbed to the disease and captured both parenchymal and non-parenchymal cell populations in epithelial tissues at high fidelity with single nucleus RNA-seq (snRNA-seq). While we have investigated the COVID-19 pathobiology of the acute respiratory distress syndrome (ARDS) lung in depth, including by spatial -omics *in situ*, the impact in other organs, including the liver, have not yet been deeply explored.

Multiple factors may underlie the COVID-19 liver phenotype, including the impact of direct infection given the expression of SARS-CoV-2 entry factors in major hepatic cell classes ^3,8,9^, systemic inflammation, drug-induced injury, and hypoxia ^3,10^. Some studies suggest the presence of subclinical liver damage, especially in the liver vasculature ^11^, with short- and potentially long-term implications.

Metabolic, vascular, and biliary alterations in COVID-19 patients could result from direct or indirect viral damage to the liver ^3^, while it was recently shown through bulk RNA sequencing and proteomics that bulk gene and protein profiles of livers identified as positive with SARS-CoV-2 present similarities to the signatures associated with multiple other viral infections of the human liver^4^. This further increases the importance of identifying its effects on infected cells and their interactions with their microenvironment. The spatial manifestation of COVID-19 phenotypes in the liver could especially be of interest due to its distinct architecture. The liver is organized in the hexagonal-shaped repeating anatomical units of the liver lobules, radiating into spatially distinct lobular zones that span from the portal triad to the central vein. The oxygen and nutrition gradients between the portal and central vein dictate liver development and define cellular function. While cellular expression programs are affected by zonation in both health and disease ^12,13^, most spatial and zonation information to date has been derived from selected markers or by concordance with animal models ^12^.

Here, we created an integrated liver COVID-19 atlas of 80,808 snRNA-seq profiles from liver samples collected at autopsy from 17 patients who succumbed to severe COVID-19, as well as whole transcriptome spatial profiling of 62 regions of interest (ROIs) from four concordant livers. By comparison with healthy controls (n=4), we generated a high-resolution map of the cellular landscape of the COVID-19 liver as well as determination of the viral impact on cell subsets, their activation states, and cell-cell communication. We used these to assess clinically relevant changes in hepatocytes and hepatic non-parenchymal cells in response to viral infection.

### A liver cell and spatial atlas in severe COVID-19

To construct a COVID-19 liver atlas, we leveraged an autopsy cohort of 17 COVID-19 patients (6 males, 11 females, ages from 30-35 to >89 years) across four medical centers from the Northeastern United States (**Table 1, Fig. 1a**) ^6,7,14^. All samples were obtained postmortem using either ultrasound-guided needle biopsy or surgical dissection by following stringent protocols established previously ^6^ (**Methods**). Most patients had multiorgan failure at the time of death. While liver function serum markers within 24 hour of death showed varying degrees of transaminitis, no patient had clinical or laboratory signs of liver failure or acute liver injury (**Extended Data Table 1**).

**Figure 1:**
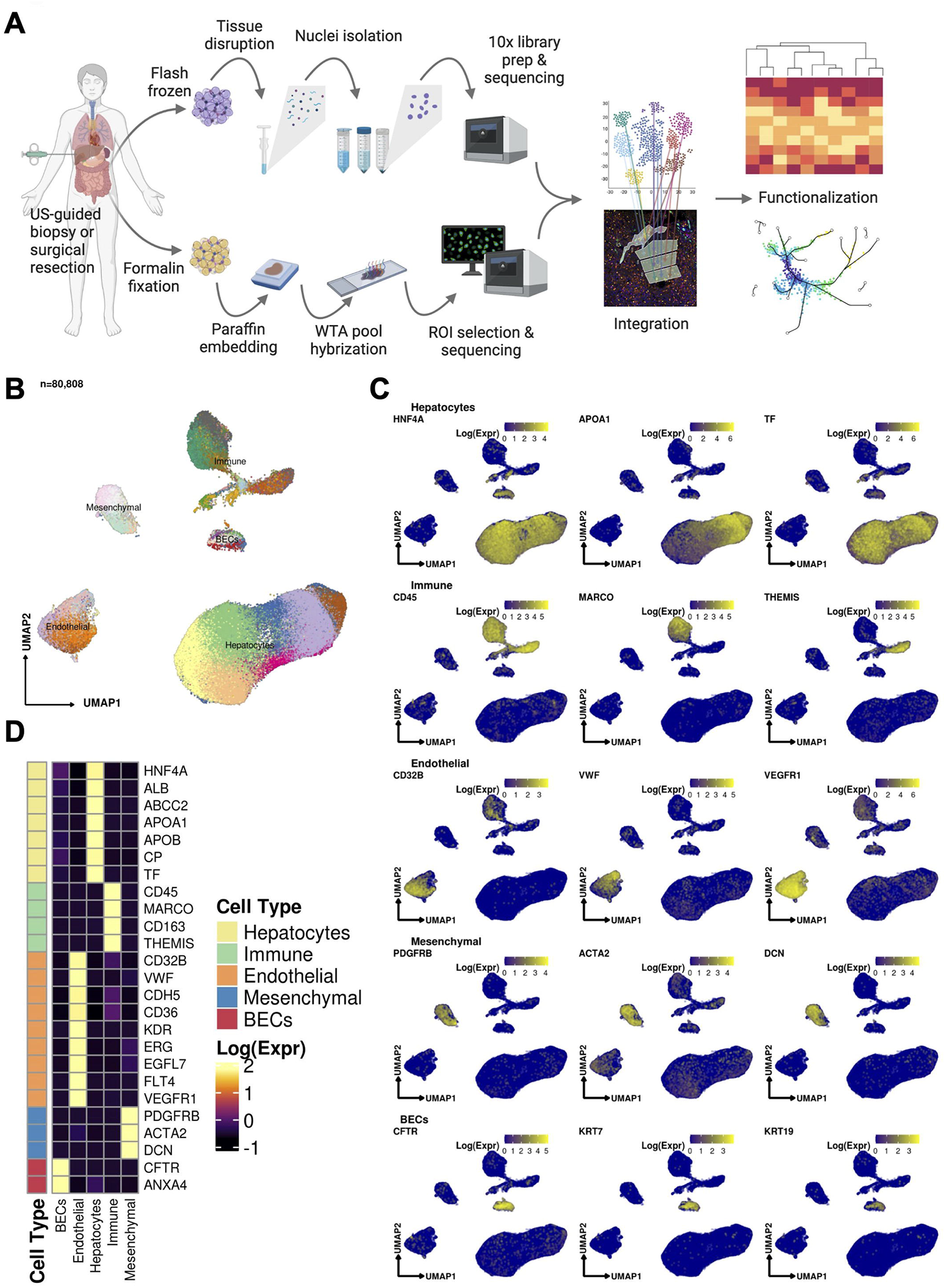
(A) Sample processing pipeline depicting sample acquisition, preparation for snRNAseq and spatial transcriptomic profiling, data generation, integration, and *in silico* functionalization. (B) Uniform manifold approximation and projection (UMAP) for all cells passing quality control (n=80,808, Hepatocytes, n = 51,605; Immune / blood cells, n = 12,346; Endothelial cells: n = 9,278, Mesenchymal cells (n = 4,647); Biliary epithelial cells / Cholangiocytes, n = 2,932). (C) Heatmap capturing the expression of marker genes across the 5 major compartments. (D) UMAP plots depicting gene marker expression for each compartment.

We used snRNA-seq to collect 80,808 high quality profiles from 17 COVID-19 patient autopsies (**Methods**) and integrated them computationally with snRNA-seq profiles from four healthy controls, prepared using a comparable protocol ^15^. Following ambient RNA removal, quality control (QC), and preprocessing (**Methods**), we implemented a batch correction pipeline to generate corrected unique molecular identifier (UMI) counts per cell ^16–18^, which facilitated marker detection and cell type identification (**Methods**). The COVID-19 nucleus profiles were partitioned into five major compartments: hepatocytes (*k* = 51,605 cells; 63.8% of all nuclei); immune/blood (*k*=12,346; 15.3%); endothelial (*k*=9,278; 11.5%), mesenchymal (*k*=4,647; 5.8%), and biliary epithelial cells (BECs) (*k*=2,932; 3.6%) (**Fig. 1b-d, Extended Data Fig. 1a,b**), spanning 50 cell subsets in distinct clusters (Cluster Dictionary provided in the **Supplementary Note**).

In parallel, we generated a spatial transcriptomic atlas from 62 Regions of Interest (ROIs) from lobular zones 1, 2, and 3, and the portal triad across four patient autopsies using the NanoString GeoMx Digital Spatial Profiling (DSP) Whole Transcriptome Atlas (WTA) platform (**Fig. 1a, Methods**). We first performed multiplexed immunofluorescence (Pan-cytokeratin (PanCK), CD45, CD68, Syto 83 (Thermo Fisher Scientific)) on the same slides to define the lobular structure by identifying the portal triad and central vein as landmarks, as well as RNA *in situ* hybridization (RNA ISH) performed on a serial section against ACE2, TMPRSS2, and SARS-CoV-2 RNA to take also into account localized viral presence (**Fig. 2a, Extended Data Fig. 1c, Methods**). We then selected 62 ROIs, corresponding to lobular zones 1, 2, and 3, and the portal triad, by the consensus opinion of an expert panel of pathologists (J.H., S.R.), hepatologists (Z.G.J., Y.P., G.S.), and technology specialists (L.P., Y.L., Y.P-J., L.T., I.S.V.). We captured the expression of over 18,300 genes on the WTA, including 27 SARS-CoV-2-relevant probes (**Extended Data Table 2**). We further developed and applied an optimized pipeline for Nanostring DSP WTA data normalization and preprocessing (**Methods**). The snRNA-seq and spatial profiles were interpreted and integrated using batch-corrected markers, a streamlined method for assigning pathway activity scores (PAS) (**Methods**), and by spatial registration of snRNA-seq profiles and signatures to decipher the localized interactions of cell types in the context of liver architecture (**Fig. 2b, Methods**).

**Figure 2:**
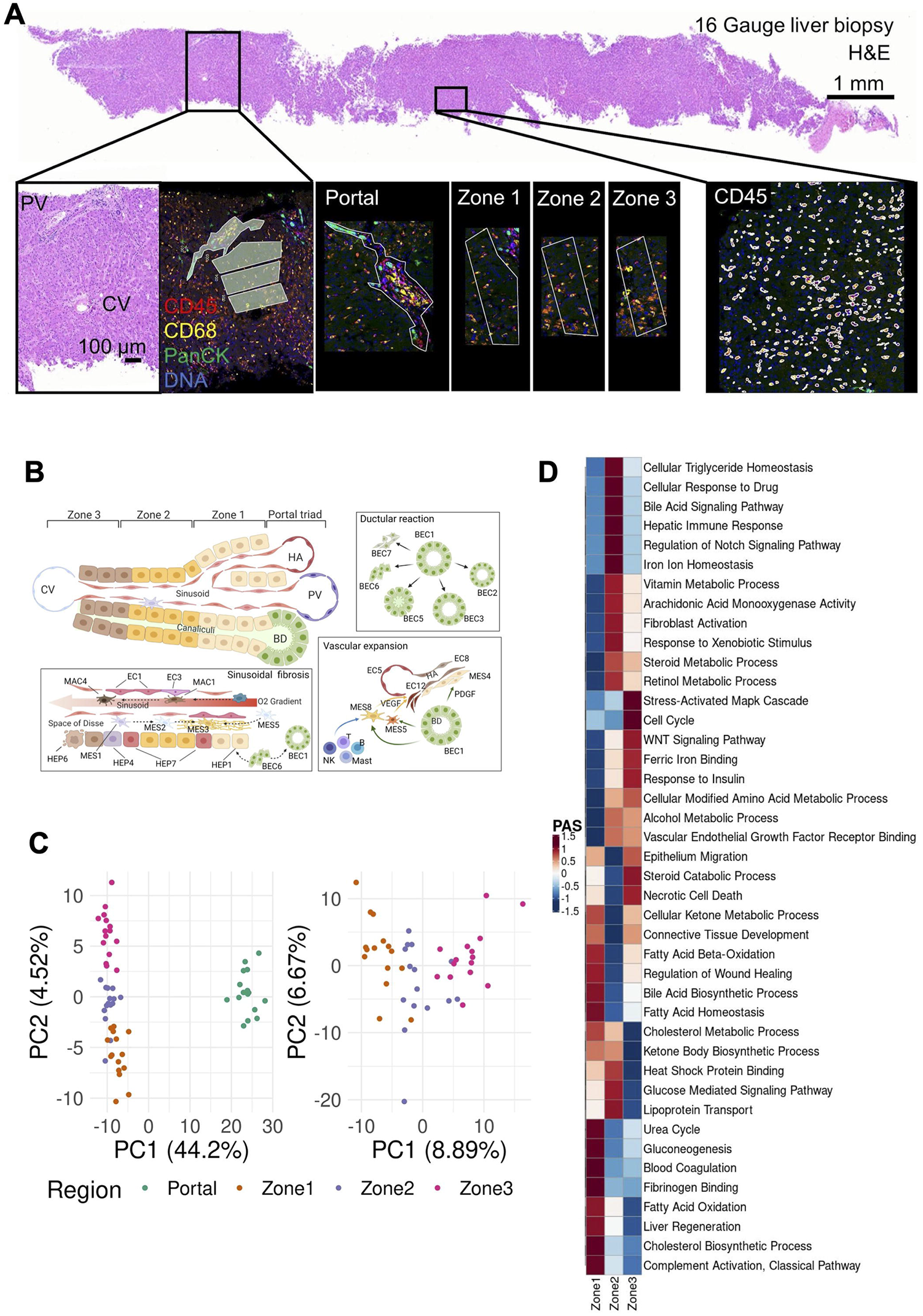
Overview of the Digital Spatial Profiling (A) Regions of Interest (ROIs), corresponding to the liver lobule and the portal area. Gene expression in each region was profiled using the NanoString GeoMx Digital Spatial Profiling (DSP) Whole Transcriptome Atlas (WTA) platform. (B) Diagram of the spatial arrangement of cellular subpopulations in the liver lobule and interactions in the context of COVID-19 (HA, hepatic artery; PV, portal vein; CV, central vein; BD, bile duct). (C) Principal component analysis (PCA) embeddings based on batch corrected probe counts of all ROIs (right) and for the liver lobule ROIs (left) reveal that the DSP WTA platform correctly separates the lobular region from the portal, and reveals significant expression differences between the 3 zones. (D) Normalized Pathway Activity Scores (PAS) between lobule regions. The DSP WTA is able to capture known zone-specific pathways as well as reveal perturbed pathways related to liver pathology and viral infection.

### Distinct zonal expression programs and their alterations in the COVID-19 liver

Each of the spatial transcriptomic ROI classes – three lobular zones and the portal triad – exhibited distinct expression profiles, with differential engagement of hepatic cellular pathways across the liver lobule, demonstrating the expected zonal division of hepatocellular function in the healthy liver ^12^ as well as its alteration in COVID-19. Principal Component Analyses (PCA) of the spatially defined expression profiles captured expression segregation between the portal triad and all lobular zones as well as among the three lobular zonal ROIs 1, 2, and 3 (**Fig. 2c**). Each region class was characterized by the differential expression of distinct region-specific markers and of functional gene sets ^12,19–21^ (**Fig. 2d**). Based on a pathway activity score (PAS) analysis (**Fig. 2d, Extended Data Fig. 1d,e, Methods**), Zone 1 exhibits high activity of transcriptional programs for lipid and glutathione metabolism, urea cycle, fatty acid and steroid biosynthesis, and lipoprotein assembly, all commonly associated with liver-specific functions. Zone 2 follows similar patterns, but with higher activity of triglyceride catabolism and fucose biosynthesis. In contrast, Zone 3 exhibited high activity of drug catabolism programs. These processes are concordant with our current functional understanding of the zonated liver and have implications for chronic liver diseases. For instance: (1) hepatic steatosis typically starts in Zone 3 ^22^ in metabolic dysfunction associated fatty liver disease (MAFLD) and alcohol-related liver disease likely due to the lower metabolic activity; (2) drug-induced liver injury is most significant in the pericentral area as a result of drug catabolism; (3) disease related to impaired metabolism may manifest preferentially in Zone 1; and (4) Zone 1 predilection of pediatric NAFLD may in part be driven by genetic variants impacting lipid and lipoprotein metabolism, such as *PNPLA3* ^23^.

In COVID-19, we found evidence of a spatially orchestrated COVID-19-specific liver phenotype, including hepatocyte proliferation in Zone 1 as well as hypoxia and stress response pathways in Zone 3, which has not been reported in healthy liver. The phenotype was reflected by high activity scores of specific pathways across liver zones and the portal triad (**Extended Data Fig. 1d** and **Extended Data Table 3 and 4**). Nonparenchymal cells showed distinct zonation of cellular physiology in the COVID-19 liver. For instance, among endothelial expression programs, differentiation programs were strongest in portal ROIs, programs for regulation of endothelial barrier establishment were highest in Zone 1, and endothelial cell chemotaxis in Zone 2 (**Extended Data Fig. 1d**). Among immune cells, portal ROIs exhibited high activity of monocyte activation and differentiation, as well as lymphocyte differentiation, whereas Zone 1 was characterized by Kupffer cell (KC) and natural killer (NK) cell proliferation, and lymphocyte migration and activation (**Extended Data Fig. 1d**). Among mesenchymal cells, portal ROIs had the highest activity of fibrogenic hepatic stellate cell (myofibroblast) activation, including response to platelet-derived growth factor (PDGF), fibroblast growth factor receptor (FGFR), and collagen/extracellular matrix production and organization pathways. Finally, Zone 3 exhibited the highest inflammation signals, including inflammasome activation, signaling by interleukins, response to cytokines, interferon-gamma binding, and inflammatory cell apoptotic processes (**Extended Data Fig. 1d**), which may be associated with SARS-CoV-2 infection and are not expected to be pronounced in Zone 3 in healthy livers. Thus, Zone 3 seems to be most severely affected in COVID-19.

### A spectrum of hepatocyte subsets from progenitors to functionally mature cells suggests plasticity of regeneration across hepatic zones

Hepatocytes were the most populous compartment in the COVID-19 snRNA-seq atlas (63.8%) (**Fig. 3a, Supplementary Note)** thanks to the ability of single-nucleus sequencing to capture this often underrepresented cell type in single-cell assays. Hepatocytes partitioned into seven subsets that spanned a continuum between two dichotomous ends: (1) primary essential liver functions, such as production of blood proteins, and (2) cell differentiation and replenishment, along with response to stress (**Fig. 3a, Extended Data Fig. 2a,b**). Regarding liver function, HEP2 cells (21.7% of hepatocytes) highly expressed genes encoding circulating blood proteins, including albumin, coagulation factors, and apolipoproteins (**Fig. 3b**), suggesting that only a fraction of all hepatocytes carry out conventional essential liver functions. HEP6 and HEP7 cells had similar profiles to those in the HEP2 subset but with high expression of acute phase proteins in HEP7 (*e*.*g*., *CRP, C3, C4a, SAA1*, and *FTH1*; a COVID-19 specific cluster; below) or apoptosis and cellular senescence pathways in HEP6 (**Fig. 3b, Extended Data Fig. 3a**). In contrast, cells in the HEP1, HEP3, and HEP4 subsets (**Fig. 3a**) exhibited lower levels of liver metabolic or synthetic function genes, but higher levels of cellular differentiation, wound healing, and signal transduction pathways (**Extended Data Fig. 3a,b**), such as the *HNF4A*/*HNF4B, YAP*/*TAZ, PPRA*/*B*/*G*, and GHR signaling pathways. HEP4 cells also expressed collagen-modifying enzymes (*P4HA1, PLOD2*; **Fig. 3b**) and pro-angiogenic factor *VEGF-A*, indicating potential regulation of hepatocyte-endothelial cell interactions. Overall, the human liver demonstrates a balance between metabolic and proliferative dynamics, as also reported in mouse liver regeneration models ^24^.

**Figure 3:**
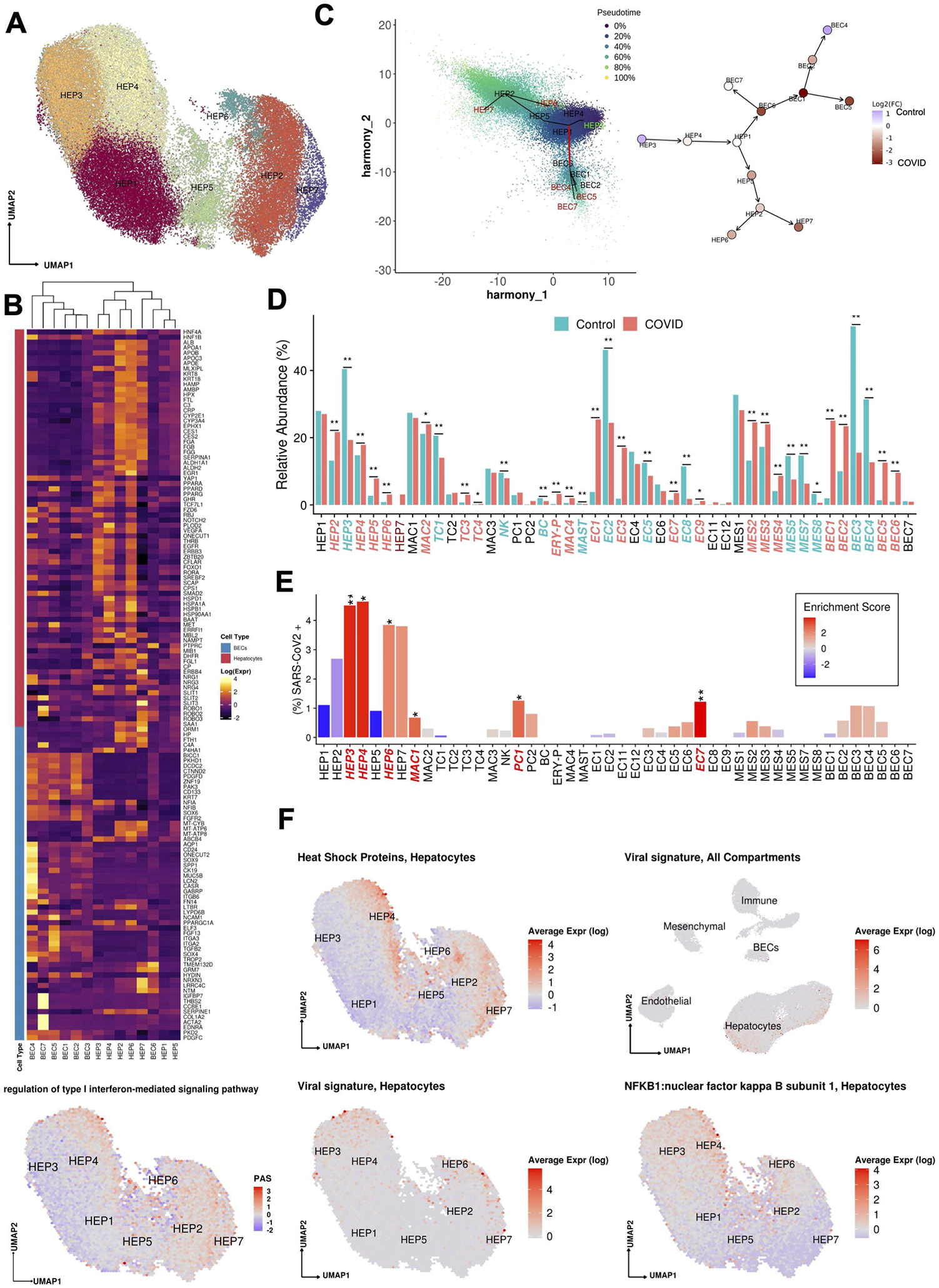
(A) Uniform manifold approximation and projection (UMAP) for Hepatocytes (HEP1 n=13,951, HEP2 n=11,187,HEP3 n=9,956, HEP4 n=9,241, HEP5 n=4,056, HEP6 n=1,612, HEP7 n=1,602). (B) Heatmap capturing the expression of marker genes across the hepatocyte and the biliary epithelial cell compartments. (C) Slingshot pseudotime values (left) projected on the 2 primary harmony embeddings across 5 lineages for hepatocyte and biliary epithelial cells from COVID-19 and healthy liver nuclei. The starting and ending lineage points are represented with green and red, respectively. Slingshot-derived lineages (right), coupled with cell composition fold-change differences between healthy and COVID-19 liver samples on a log2 scale. (D) Cell proportion differences between COVID-19 and healthy liver samples. Significantly different proportions are marked in red (higher in COVID-19), in blue (higher in Controls), and denoted with *(* FDR < 0.05, ** FDR < 0.01; Binomial Generalized Linear Mixed Model). COVID-19-specific clusters are denoted with dark red. (E) Abundance of SARS-CoV-2 RNA+ nuclei in the snRNAseq clusters. The bars are colored by the scaled viral enrichment score estimated per cluster. Significantly enriched clusters are marked in red and denoted with * (* FDR < 0.05, ** FDR < 0.01; Viral enrichment test). (F) Uniform manifold approximation and projection (UMAP) plots depicting the average expression of different heat shock proteins (HSPA1A, HSPA1B, HSPA5, HSPA6, HSPA9, HSPB1, HSPD1) in Hepatocytes (upper left), pathway activity scores for GO term “regulation of type I interferon-mediated signaling pathway” (GO:0060338, bottom left), the viral load in all the cellular compartments (upper right), in Hepatocytes (lower middle), and the average expression on NFKB1 in Hepatocytes (lower right).

Trajectory analysis of epithelial cells (hepatocytes and cholangiocytes) from both healthy and COVID-19 livers (**Methods, Fig. 3c**) suggests a differentiation path from HEP3 cells, a cell population with the highest pathway activities related to cell replication and expressing *WNT* and *NOTCH* signaling pathway genes (*e*.*g*., *TCF7L1, TCF7L2, FZD6, RBPJ, NOTCH2*; ^25^) to the highly differentiated HEP2 cells, through HEP4, 1, and 5 intermediates, with HEP6 and HEP7 cell populations directly derived from HEP2. The hepatocyte population is known to be maintained both through mitosis of mature hepatocytes and differentiation from hepatic progenitor cells (HPCs) ^26^. As HPCs give rise to both BECs and hepatocytes ^27^, and injured hepatocytes can transition into HPCs ^28^, we included both epithelial (hepatocyte and BEC) compartments, finding that HEP1 cells were an intermediate across hepatocytes and cholangiocytes (**Fig. 3c**). BEC differentiation trajectories are further discussed below.

### Hepatocyte composition and differentiation are altered in COVID-19

Contrasting healthy and COVID-19 cellular landscapes (**Methods, Extended Data Table 5**) reveals extensive remodeling of the hepatocyte compartment in COVID-19 (**Fig. 3d**), as well as the emergence from HEP2 cells of a COVID-19-specific HEP7 cluster, expressing acute phase proteins (**Fig. 3b-d**). Consistent with a model of regenerative capacity loss in COVID-19 liver, the proportion of the less differentiated HEP3 cells was reduced (FDR=3.63×10^−54^, OR=0.352, Binomial GLMM) whereas proportions of HEP2, HEP4, HEP5, and HEP6 cells were identified as increased (HEP2,4,5,6: FDR=8.50×10^−26^, 2.37×10^−6^, 8.60×10^−6^, 2.22×10^−48^; OR=1.82, 1.26, 3.04, 3.52; Binomial GLMM; respectively) or only present in COVID-19 samples in the case of HEP7 **(Fig. 3c,d**). Comparing the COVID-19 specific HEP7 cells to the closely related HEP6 cells, shows an inverse CEBPA/CEBPB ratio, demonstrating a metabolic vs. acute phase regulation expression program ^29^. Notably, HEP4 hepatocytes also exhibit low *HNF4A, APOB*, and high *SCARB1, STAT3*, and *HIF1A*, a phenotype identified using bulk proteomics on severe COVID-19 patient livers, and hypothesized to be driven by the combination of hypoxia and activation of *STAT3*, leading to a reduction of the differentiated hepatocyte pathways orchestrated by down-regulation of *HNF4A* ^30^. The trajectory analysis reveals not only a reduction of lineages concordant to the differential cellular proportions observed, such as the increase of cells in the stressed HEP4 state, but also COVID-19-specific lineages, with high proportion of cells in the terminally differentiated HEP2 state and in the COVID-19-specific acute response HEP7 cluster (**Fig. 3c, Extended Data Fig. 3c,d)**.

### SARS-CoV-2 RNA+ cells are enriched in hepatocyte subsets and associated with specific expression changes

We analyzed the donor and cell type specific distribution of SARS-CoV-2 sequencing reads to determine the presence of viral transcripts in liver cells. Specifically, we called each nucleus profile as SARS-CoV-2 RNA+ or SARS-CoV-2 RNA-by comparing the observed viral unique molecular identifier (UMI) counts to the ambient pool (a potential source of viral RNA contamination) and then tested for the enrichment of SARS-CoV-2+ nuclei in each cell type (**Methods)**. Hepatocytes were the most enriched for SARS-CoV-2 RNA+ nuclei, particularly within the least differentiated (HEP3, 4: FDR = 1×10^−8^, 1×10^−8^; ViralEnrichment test; respectively) and most differentiated clusters (HEP6, HEP7: FDR=0.040, 0.066; ViralEnrichment test; respectively) (**Fig. 3e**).

Viral RNA levels were positively associated with the expression of multiple heat shock proteins (*HSPA1A, HSPA1B, HSPA5, HSPA6, HSPA9, HSPB1, HSPD1*), which were highest in HEP3, 4, 6, 7 (**Fig. 3f, Fig 4a,b**), suggesting activation of unfolded protein response to cellular stress in these subsets. In HEP4, profiles with higher viral UMIs also exhibited high *NF-kB* expression (**Fig. 3f**) suggesting an activation of an inflammatory response, concomitant with epithelial cell SARS-CoV-2 infection^31^. Infected cells also overlapped with high pathway activity score (**Fig. 3f, Methods)** for the gene ontology (GO) term “regulation of type I interferon-mediated signaling pathway” (GO:0060338). Interferon signaling pathways were identified as enriched in a bulk RNA-seq analysis of 5 samples from SARS-CoV-2 positive livers, as characterized by PCR, when compared against 5 SARS-CoV-2 negative liver samples^4^.

**Figure 4:**
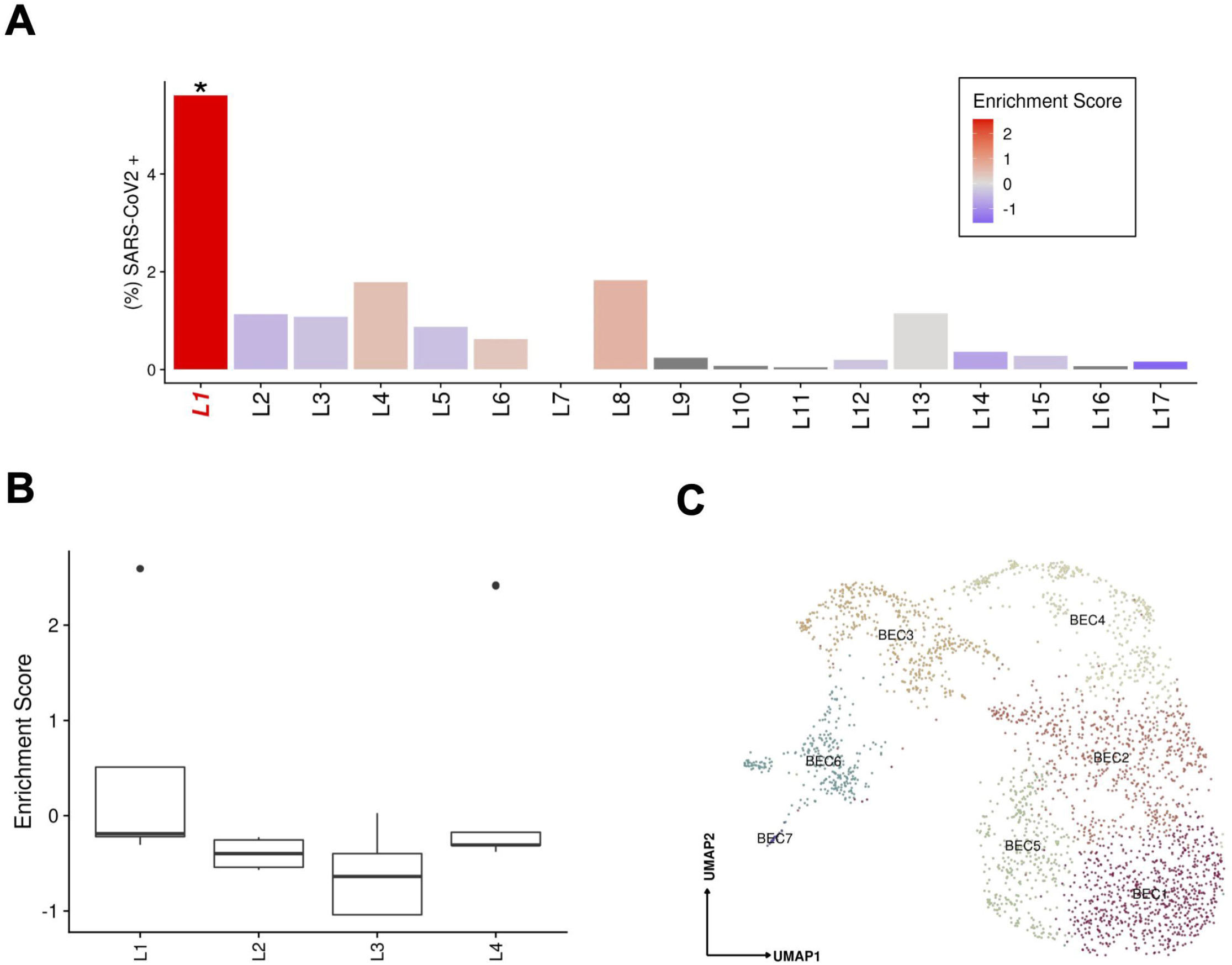
(A) Abundance of SARS-CoV-2 RNA+ nuclei in the snRNAseq data for each donor. The bars are colored by the scaled viral enrichment score estimated per donor. Only donor L1 has a significant viral enrichment score (* FDR < 0.01; Viral enrichment test). (B) Distribution of the NanoString GeoMx DSP SARS-CoV-2 probe enrichment score across donors. Donor L1 has a significantly higher enrichment score (FDR = 0.037, t-test) compared to the rest of the donors (L2-4). (C) Uniform manifold approximation and projection (UMAP) for Biliary epithelial cells (BEC1 n=736; BEC2 n=687; BEC3 n=457; BEC4 n=373; BEC5 n=371; BEC6 n=281; BEC7 n=27).

SARS-CoV-2 RNA+ cells and viral UMIs also varied across patients. Donor L1 cells were significantly (FDR<0.01, viral enrichment score) enriched for SARS-CoV-2 RNA+ nuclei (9-fold higher proportion of enriched nuclei vs. average of all other donors) (**Fig. 4a**). Since the ability to detect viral UMIs can be affected by the total number of UMI counts and the number of genes detected (**Extended Data Fig. 4**), we also tested for enrichment in SARS-CoV-2 viral-specific probes in the extended NanoString GeoMx DSP WTA assay (**Methods)**. Donor L1 has a significantly higher enrichment score (FDR=0.037, t-test) for viral probe counts compared to the other donors (**Fig. 4b**). The significant enrichment in donor L1 for SARS-CoV-2 RNA in both the snRNA-seq and GeoMx DSP assays was consistent with the viral abundance estimated by quantitative RT-PCR using liver tissue from the same samples (**Extended Data Table 6**). Interestingly, the higher viral load detected by snRNA-seq, GeoMx, RT-PCR, and RNA ISH (**Extended Data Fig. 1c**) was not associated with gross abnormality of the liver tissue by conventional H&E staining. Consistent with previous reports (Delorey et al. 2021; Wölfel et al. 2020; Walsh et al. 2020), we found a negative, but not statistically significant correlation between the duration from symptom start to death, and the enrichment score for SARS-CoV-2 (p=0.2852, Spearman ρ= -0.336) (**Extended Data Table 7**).

### Pathological expansion of the cholangiocyte compartment in COVID-19

BECs (3.6% of COVID-19 patient liver nuclei, **Supplementary Note**) expressed the lineage markers *CFTR, KRT7*, and *KRT19*, and spanned a broad spectrum, partitioning to six main subsets (**Fig. 4c**): two subsets of differentiated cholangiocytes (BEC1, 2), three of reactive cholangiocytes/HPCs (BEC4,5,6), and one minor subset of cholangiocyte with mesenchymal features (BEC7). BEC3 expressed highly MT genes and hepatocyte-specific markers *CPS1, ALB, HNF4A, C3, ABCB4*, which could potentially be doublets. BEC1 and 2 were closely related fully differentiated small cholangiocytes lining small caliber bile ducts ^32^, expressing secretin receptor *SCTR, BCL2*, and primary cilia genes (*e*.*g*., *BICC1, PKHD1, DCDC2, CTNND2, PKD2*, but not *CYP2E1*; **Fig. 3b**), while BEC1 expressed lower levels of *PDGFD, ZNF19, PAK3, ONECUT1*, and CD133 compared to BEC2.

BEC4, 5, and 6 subsets each had a distinctive profile, consistent with either “reactive” cholangiocytes/hepatic progenitor cells (HPCs) or with a pro-fibrogenic “ductular reaction” in chronic liver diseases (Roskams and Desmet 1998). BEC4 cells comprised osteopontin-positive reactive cholangiocytes/hepatic progenitor-like cells (HPCs), expressing *SPP1, SOX9* ^33^, *LYPD6, CASR, HNF1B, ONECUT1*/*2*, and *GABRP*, as well as progenitor cell response genes (*ITGB6, FN14*/*TNFRSF12A, LTBR*). BEC5 were NCAM1^+^ immature, reactive cholangiocyte/HPCs ^34^, co-expressing *ITGA2*, progenitor cell markers (*SOX4, CK19, TROP2*, CD133), and potent pro-fibrogenic mediators (*FGF13, PPARD, PDGFC*, and *TGFB2*). BEC6 were a neuroendocrine subset of cholangiocytes ^35^, expressing neural markers (*TMEM132D, GRM7, HYDIN, NRXN3, LRRC4C, NTM*). Trajectory analysis suggests that BEC6 cells may form a potential transition state between hepatocytes and cholangiocytes (**Fig. 3c**), consistent with previous findings ^27^. BEC7 comprised a minor subset of activated cholangiocytes co-expressing both epithelial and mesenchymal genes (*IGFBP7, THBS2, CCBE1, COL1A2, ACTA2, EDNRA*) and many cell-cell communication genes, especially with the endothelial compartment (*FGF, PDGF, VEGF* ligands/receptors) (**Extended Data Fig. 5a-c**), and is connected to BEC6 in the trajectory analysis **(Fig. 3c)**.

Compared to normal liver (**Fig. 3c,d**), BEC4 (and BEC3s) were reduced (FDR=2.36×10^−6^, 1.32×10^−18^; OR=0.318, 0.162; Binomial GLMM; for BEC 4,3. respectively) and BEC1, 2, 5, and 6 increased in COVID-19 liver samples (BEC1,2,5,6: FDR=3.80×10^−15^, 2.22×10E^-5^, 7.74×10^−10^, 2.21×10^−6^, respectively; OR=16.577, 2.736, 10.413, 11.482, respectively; Binomial GLMM), showing an extensive pathological restructuring of the cholangiocyte compartment. Spatial transcriptomics revealed that while BEC1,2 and 4 signatures mapped to portal tracts as expected (**Extended Data Fig. 6a**), HPC-like BEC6 and 7 had mixed lobular and portal distribution in COVID-19 liver, consistent with pathological “ductular reaction” expansion into the hepatic lobule ^36^. We validated this observation by CK19 staining in these livers, which revealed a presence of ductular reaction in all samples, ranging from minimal to extensive multifocal ductular proliferation extending well into the liver lobule (Donors L1-4, **Extended Data Fig. 6b**).

### Kupffer cell proliferation and emergence of an erythrocyte progenitor population in COVID-19

The immune and blood cell compartment of COVID-19 livers (15.3% of COVID-19 patient liver nuclei) spanned monocytes/macrophages/Kupffer (KCs), T cells, B cells, natural killer (NK) cells, and mast cells in diverse cellular states (**Fig. 5a, Supplementary Note**).

**Figure 5:**
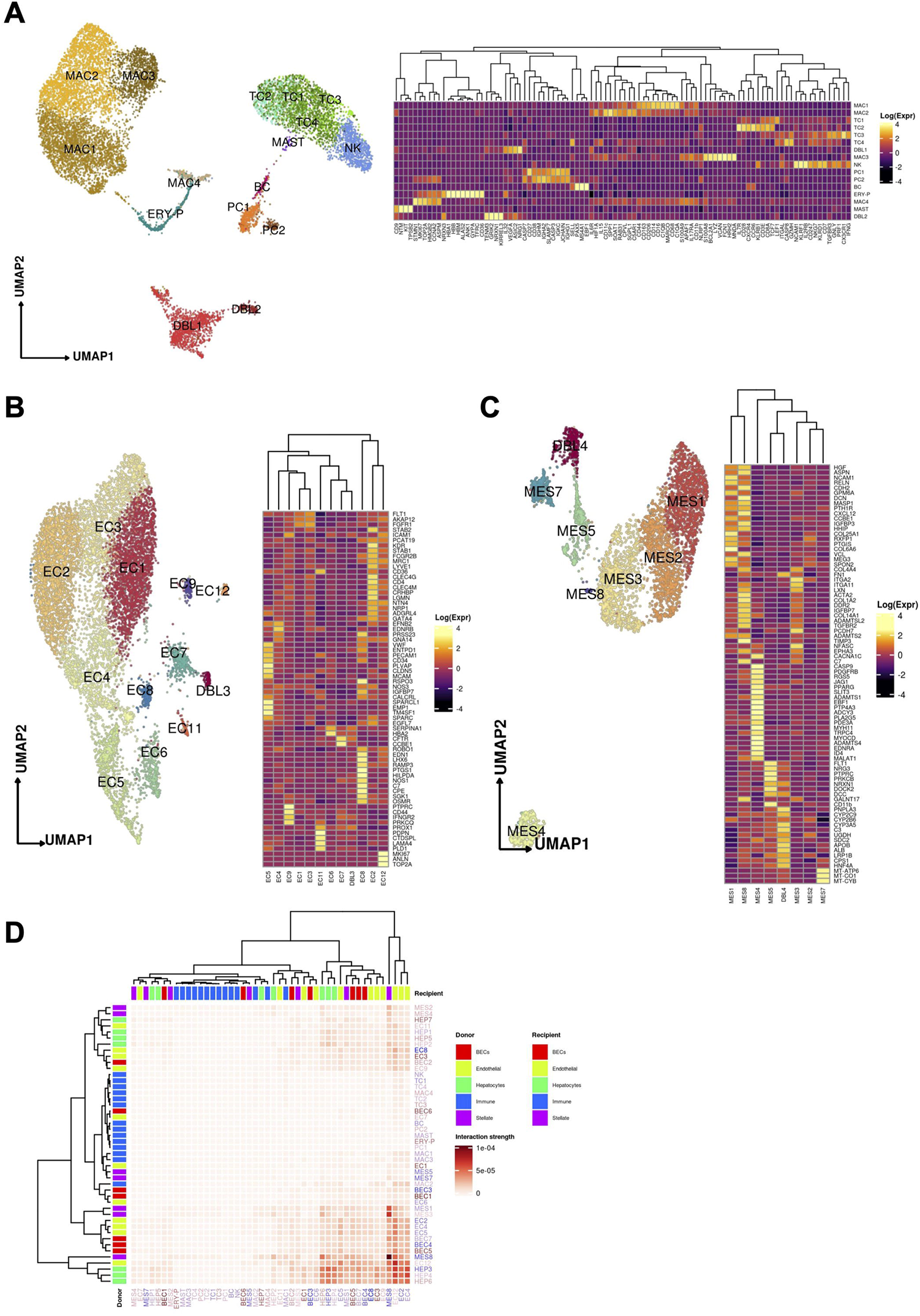
(A) Uniform manifold approximation and projection (UMAP) for the (A) Immune / blood, (B) Endothelial cell, and (C) Mesenchymal cell compartments ((A) *Immune:* MAC1 n = 2,798, MAC2 n = 2,601, TC1 n = 1,522, TC2 n = 388, TC3 n = 327, TC4 n = 29, DBL1 n = 1,331, MAC3 n = 1,038, NK n = 857, PC1 n = 397, PC2 n = 124, BC n = 124, ERY-P n = 359 MAC4 n = 222, MAST n = 36 DBL2 n = 193; (B) *Endothelial:* EC1 n = 2,338, EC2 n = 2,247, EC3 n = 1,563, EC4 n = 1,117, EC5, n = 795, EC6 n = 379, EC7 n = 328, EC8 n = 166, EC9 n = 116, DBL3 n = 91, EC11 n = 73, EC12 n = 65, (C) *Mesenchymal:* MES1 n=1,223, MES2 n=1,065, MES3 n=1,040, MES4 n=374, MES5 n=328, MES6 n=312, MES7 n=275, MES8 n=30). Heatmaps capturing the expression of marker genes across the 3 distinct major compartments are displayed. (D) Heatmap portraying the cell-cell communications between the cell populations. The color gradient indicates the strength of interaction between any two cell groups. Recipient/Donor cell-type color is portrayed in a blue (healthy) to red (COVID-19) gradient, relevant to the cell composition fold-change differences between healthy and COVID-19 liver samples.

Both the myeloid and T cell compartments were remodeled in the COVID-19 liver compared to healthy controls (**Fig. 3d**). Naive CD8+ T cells with high expression of *LEF1* and *TCF7* (TC1) were significantly decreased in COVID-19 liver (FDR=1.45×10^−9^, OR=0.629, Binomial GLMM), while cytotoxic effector/memory T cells (TC3), expressing *IFNγ, CX3CR1, TGFBR3, GNLY*, and *GZMH*, and the apoptotic naive T cell-like (TC4) population were both significantly increased in the COVID-19 liver (FDR=1.69×10^−4^, 2.59×10^−2^; OR=4.127,1.969 Binomial GLMM). In the myeloid compartment, there were no differences in classical Kupffer cells (KCs) (MAC1) or inflammatory KCs (MAC3) (MAC1, MAC3: FDR=0.231, 0.154; OR=0.925, 0.873; Binomial GLMM, respectively), but an increased proportion of MAC2 cells was observed in COVID-19 (FDR=1.86×10^−2^, OR=1.182, Binomial GLMM), an intermediate phagocytic macrophage phenotype with lower expression of *MARCO* and CD164 but increased expression of phagocytic markers (*C5AR1, CPVL*, CD206). None of the macrophage subsets expressed high levels of chemokine receptors (*CCR2, CCR5, CXCR3*), indicating a deficiency of infiltrative monocyte derived macrophages, which potentially reflects a degree of immune exhaustion and/or pulmonary tropism.

The atlas also captured several proliferating cell populations that have not been previously identified in human liver single-cell studies, were nearly-exclusive to COVID-19 samples, and may play important roles in regeneration. In particular, a small subset of proliferating Kupffer cells (MAC4), were significantly increased in COVID-19 livers (FDR=7.19×10^−4^, OR=3.395, Binomial GLMM) (**Fig. 3d**). Kupffer cells can replicate following tissue injury and were recently reported as the first cell type to enter a proliferating program in mouse liver regeneration ^37^, but have not been until now reported in human samples. MAC4s clearly recapitulate the scRNAseq signature identified in mouse liver samples following injury (**Extended Data Fig. 6c, Methods**). Moreover, erythrocyte precursors (ERY-P) were detected almost exclusively in the COVID-19 liver (FDR=2.37×10^−6^, OR=12.554, Binomial GLMM), expressing a combination of hemoglobin and glycophorin genes, proliferation genes, and additional genes not present in mature red blood cells, such as CD71/*TFRC*, which are rarely encountered outside the bone marrow in adults. These cells may be responsible for extramedullary hematopoiesis in the setting of hypoxia, modulate immune response in virus infection, and participate in hepatogenesis as shown in fetal liver ^38,39^.

### Disrupted zonation and differentiation of endothelial cells in COVID-19

Cells in the endothelial compartment (11.5% of COVID-19 patient nuclei) spanned 12 subsets, including liver sinusoidal endothelial cells (LSEC) and other endothelial cell (EC) populations in an 8:1 ratio (**Fig. 5b, Supplementary Note**).

Endothelial cell composition was substantially impacted in COVID-19 *vs*. healthy liver (**Fig. 3d**). EC1 cells, the largest endothelial subset in COVID-19 liver samples, were significantly increased in proportion compared to healthy liver (FDR=2.76×10^−23^, OR=8.63, Binomial GLMM). These cells expressed *VEGFR1, FGFR1*, and A-kinase Anchoring Protein 12 (*AKAP12*), but were *VEGFR2* negative. *FGFR1* is upregulated in cholestatic liver injury in mice, which provokes maladaptive fibrogenesis ^40^, while *AKAP12* deficiency is linked to VEGF-induced endothelial cell migration ^41^, regulates cell adhesion ^42^, and supports the integrity of the blood brain barrier during ischemic injury ^43^. In the liver, *AKAP12* also modulates the activity of hepatic stellate cells (HSC) in liver injury ^44^. Thus, EC1 represents a LSEC-derived profibrotic niche in response to systemic illness, either directly or indirectly from SARS-CoV-2. Conversely, EC2s, typical liver sinusoidal endothelial cells (LSEC) with high lymphatic vessel endothelial hyaluronan receptor (*LYVE1*) expression, and EC8s with features of classical vascular endothelial cells and high anti-inflammatory gene *C7* expression ^45^ were both significantly reduced in the COVID-19 samples (EC2, EC8: FDR=7.10×10^−11^, 5.16×10^−29^; OR=0.378,0.142; Binomial GLMM; respectively) (**Fig. 3d**). EC3s likely represented transitional states from EC2 to EC1 and were also increased in COVID-19 livers (FDR=2.91×10^−19^, OR=10.571, Binomial GLMM).

Notably, two clusters of rare cell populations were detected almost exclusively in COVID-19 livers, which may partly reflect the larger number of profiled nuclei. EC11 cells, a rare subset of *FLT1* (*VEGFR1*) negative cells (0.8% of endothelial cell nuclei; 0.09% of all profiled nuclei; FDR=7.61×10^−1^, OR=9.665, Binomial GLMM) are lymphatic endothelial cells, which are potentially captured in our COVID dataset due to the larger number of profiled nuclei. Another rare subset detected primarily in COVID-19 liver were EC12 cells (FDR=1.76×10^−1^, OR=2.864, Binomial GLMM), expressing proliferation and angiogenesis-associated genes. This subset is reminiscent of replicating endothelial cells observed in mouse lung following influenza injury ^46^. Using pathway activity scores, EC12 cells clearly recapitulated the cell signature observed in influenza infected mice (**Extended Data Fig. 6d, Methods**).

### Fibrogenic activation in the mesenchymal compartment in COVID-19 patient livers

The eight subsets of mesenchymal cells (5.8% of COVID-19 nuclei) represented all major cell lineages found in the liver, including quiescent and activated hepatic stellate cells (HSCs), smooth muscle cells (SMCs), myofibroblasts (MFs), and fibrocytes (**Supplementary Note**).

Mesenchymal cell proportions shifted substantially in COVID-19 liver, consistent with profibrotic HSC activation (**Fig. 5c and 3d**). While the proportions of quiescent HSCs (qHSCs, MES1) – the largest mesenchymal subset – were unchanged between healthy and COVID-19 livers (FDR=0.121, OR=0.807, Binomial GLMM), partially activated HSCs (aHSCs) (MES2) and extracellular matrix (ECM)-associated HSCs (MES3) were both significantly increased in COVID-19 livers (MES2, MES3: FDR=1.44×10^−8^, 9.21×10^−4^; OR=2.149, 1.508; Binomial GLMM; respectively), as were smooth muscle cells (SMCs) (MES4) (FDR=1.66×10^−4^, OR=2.181, Binomial GLMM). Conversely, both putative bone-marrow-derived fibrocytes ^47^ (MES5) and a minor subset of activated myofibroblasts (MES8) were decreased in proportion in COVID-19 *vs*. healthy liver (MES5, MES8: FDR=3.28×10^−9^, 1.09×10^−2^; OR=0.479, 0.205; Binomial GLMM; respectively). MES7 cells exhibited high expression of mitochondrial genes and low nuclear mRNA counts pointing to apoptotic cells or a technical artifact.

As expected, both MES1 (quiescent HSCs) and MES2 (activated HSCs) demonstrated translobular localization in the spatial analysis (**Extended Data Fig. 6a**), indicative of *in situ* activation of perisinusoidal qHSCs in response to parenchymal injury. Importantly, HSC activation was validated by immunohistochemistry for the classical HSC activation marker alpha-SMA, demonstrating a massive fibrogenic activation of HSCs across all studied livers (**Extended Data Fig. 6b, Methods**). In contrast, MES3 (ECM-associated HSCs), MES4 (SMCs), MES5 (fibrocytes), and MES8 (activated myofibroblasts), were mapped to the portal tract (**Extended Data Fig. 6a**). Surprisingly, we were not able to identify portal fibroblasts (PF) in the mesenchymal compartment based on PF-specific markers reported in the literature ^48,49^. This is consistent with evidence that collagen-producing myofibroblasts are a progeny of pericyte-like qHSCs, as suggested in fate-tracing studies in mice ^50^, and does not appear to support the appreciable contribution of PFs ^48,49^ to the pool of fibrogenic effector cells in the human liver in the setting of subacute liver injury.

### Cellular communication networks reveal active fibrogenesis mediating altered cellular programs in COVID-19

Cell-cell communication analysis in COVID-19 donor snRNA-seq data (**Methods**) revealed a potential multi-cellular hub of interacting mesenchymal cells, endothelial cells, and hepatocytes (**Fig. 5d, Extended Data Fig. 5a**). The hepatocyte and endothelial compartments demonstrated signaling through the *ERBB* family of proteins, including neuregulin (*NRG*) and epidermal growth factor (*EGF*), as well as the *TGF-*β family of proteins, including the central pro-fibrogenic cytokine transforming growth factor beta (*TGF-*β*1*), and bone morphogenetic protein 5 and 6 (*BMP-5, -6*) (**Extended Data Fig. 5b,c**). This finding is consistent with their previously reported role in liver tissue regeneration, cellular homeostasis, and extracellular matrix remodeling, associated with scarring ^51–56^.

We identified a robust *VEGF* signaling network that predominantly emanates from the hepatocyte compartment. The high contribution by the *VEGF-A* ligand correlates with its reported upregulation under hypoxic conditions and its role in maintenance of LSEC differentiation and of liver regeneration by enhancing liver endothelial cell communication with neighboring parenchymal cells ^56–59^. The *LIGHT* and *CXCL* signaling networks presented a distinguishable narrow number of cell-cell interactions with strong communication probability. Tumor necrosis factor superfamily 14 (TNFSF14) was the main driver of the former network with a markedly strong interaction between subclusters HEP2 and HEP5. This interaction could represent an underlying homeostatic mechanism between distinct hepatocytes responsible for regulating *TGF-*β*1* expression in liver fibrosis ^60^. Interestingly, *TGF*β-centric communication was observed between MES8 and HEP7 cells (a COVID-19-specific subset), suggesting stressed hepatocytes could be driving fibrogenic HSC activation. In addition, HEP7 also produces *CXCL12*, which promotes angiogenesis, inflammation, and has been shown to cause fibrogenesis in the lung ^61^ (**Extended Data Fig. 5b, c**). The cell-cell communication pathways support a diverse source of fibrogenic activation, involving hepatocytes, cholangiocytes, endothelial and immune cells, in contrast to an immune cell dominated framework seen in many chronic liver diseases.

### Histopathology validation of an extensive pro-fibrotic cellular phenotype of COVID-19 livers

To validate the insights from our atlas, we performed a liver histopathology survey of the four cases (Donors L1-4, **Methods**), where snRNA-seq and GeoMx assays were performed. Surprisingly, a common striking pathology feature of all four COVID-19 livers was the stellate cell activation and sinusoidal fibrosis, ranging from moderate in L1 to massive in L4. Upon further review of the medical records, none of the four donors had extensive history of primary liver disease or severe liver injury in the 72 hours prior to death. Three out of four patients also demonstrated moderate to extensive ductular reaction/cholangiocyte proliferation (**Extended Data Fig. 6b and Extended Data Table 8**). This is consistent with the increased proportion of activated/transdifferentiated mesenchymal and cholangiocytic cell subsets identified in our snRNA-seq. Although pro-fibrogenic and HSC activation pathways were observed in the cell-cell communication analysis, they cannot completely explain the great extent of HSC activation observed in the histopathological analysis. Thus, extrahepatic, systemic signals may additionally contribute to the activation of HSCs and fibrosis in the liver of severe COVID-19. Since severe COVID-19 has features of an atypical viral sepsis-like condition that goes on for an extended period of time ^62^, our findings therefore share features of the low-grade inflammation, stellate cell activation, ductular reaction, and hepatic fibrosis observed in experimental sepsis in mice ^63^.

## Discussion

We have generated a cellular and spatial atlas of the COVID-19 liver by integrating snRNA-seq and spatial transcriptomics on autopsy samples obtained from patients who died from COVID-19. We acquired >80,000 high quality single nucleus profiles with >50% hepatocyte representation, providing us with a rich, granular dataset, even for rare cell subsets.

We observed extensive pathological restructuring of the cellular and expression landscape in COVID-19 livers, suggesting hepatocellular injury, ductular reaction, neo-vascular expansion, and fibrogenesis. Based on viral RNA reads, we identified human hepatocytes infected by SARS-CoV-2 and characterized their expression profiles, while also capturing indirect and systemic effects of COVID-19 on hepatocyte populations. The highest number of SAR-CoV-2 viral RNA UMIs were found in hepatocytes, while a previously proposed cholangiocyte-tropism ^64^ in the liver was not seen. Viral RNA UMI-enriched hepatocytes exhibited high expression of acute phase and pro-inflammatory proteins, with increased heat shock protein gene expression, likely a response to unfolded proteins, secondary to viral replication; and *NF-kB* expression, consistent with the available literature for other epithelial cell types^65^. Our results also recapitulated the observation of high Interferon signaling pathway activity, as were suggested in a bulk RNA-seq analysis of 5 samples from SARS-CoV-2 positive livers compared against 5 SARS-CoV-2 negative liver samples^4^.

Meanwhile, profibrogenic/reactive cholangiocytes were identified as characteristic populations expanding in the COVID-19 liver, representing a pathological “ductular reaction” - an extensive remodeling and scarring of biliary compartment, secondary to local as well as systemic liver injury ^36^. This striking observation was validated by conventional immunohistochemistry and is consistent with emerging reports of COVID-19-induced sclerosing cholangitis (fibrotic disease of bile ducts) ^66^, which in most severe cases may require liver transplantation ^67^.

We also found extensive changes in the composition and expression programs of non-parenchymal cells across the liver lobule and portal triad in COVID-19. Endothelial cell population proportions are significantly altered in COVID-19 livers, with the emergence of a large population of *FGFR1* and *AKAP12-*positive cells that may contribute to angiogenesis and promote fibrosis ^68,69^. In the immune compartment of the COVID-19 liver, we observed KC proliferation and erythrocyte progenitors for the first time in a human single-cell study. We also observed activation of mesenchymal stellate cell/myofibroblast cells both in the liver lobule and portal areas, which were validated by immunohistochemistry staining, and an expansion of smooth muscle cell population in the COVID-19 liver samples. This pattern of fibrosis cannot be explained by underlying chronic liver disease and is likely caused by a combination of localized and systemic, sepsis-like effects of severe COVID-19 ^63^. These cellular and expression changes induced by COVID-19, despite an absence of significant tissue injury, point to subclinical yet profound effects of COVID-19 on the human liver, and may carry long-term health implications for those who recover from acute infection.

Our study captured the complexity of liver biology at high resolution, providing new insights into cellular plasticity and regeneration in the liver. Based on their RNA expression profiles, a significant proportion of the hepatocytes do not appear to contribute directly to liver function by conventional definitions, while reflecting other processes such as cellular differentiation, growth, and wound healing. Compared to previous single-cell studies, we did not observe a strict zonated distribution of hepatocyte clusters; rather, several hepatocyte subtypes intercalate in a mosaic pattern, which could be a result of augmented regeneration in response to COVID-19. Similarly, in the BEC compartment, we characterized rarely identified cells, such as neuroendocrine cholangiocytes, and a bidirectional trajectory axis between cholangiocytes and hepatocytes with specific cell transition states between these cell types, not previously reported in human samples. Other hematopoietic lineage cells were found to be in a proliferative state, including erythrocyte progenitors and plasmablasts. The former are not commonly encountered outside the bone marrow in adults, while the latter further support the recent observations made by Dominguez Conde, *et al*.^*70*^ showing the presence of this population along with ITGA8 positive plasma cells in the human liver.

Our study was limited by including a relatively small number of patients (n=17) with a severe COVID-19 phenotype, not enabling us to directly assess moderate and less severe manifestations of the disease. As all samples were analyzed early in the pandemic, they cannot inform impact from vaccination, and reflect only the very early lineages of the virus. Nevertheless, this extensive dataset offered unique insights on the sub-clinical COVID-19 liver phenotype and biology, while its very high granularity and complementary methods enable it to become the foundation of future meta-analyses and could complement basic, clinical, and translational research efforts.

## Materials and Methods

### Patient cohorts

An autopsy cohort of 17 COVID-19 patients (6 males, 11 females, ages from 30-35 to >89) was collected from 4 medical centers from the Northeastern United States during the first wave of the pandemic (**Table 1**). For all patients, consent was acquired by their healthcare proxy or next of kin prior to their inclusion to the study. Exclusion criteria included high post mortem interval (>24h) and HIV infection. All samples were obtained post mortem using either ultrasound-guided needle biopsy or surgical dissection. All sample collection procedures were reviewed by the IRB of the relevant hospital. The related protocols were: Beth Israel Deaconess Medical Center (IRB 2020P000406, 2020P000418), Brigham and Women’s Hospital and Massachusetts General Hospital (2020P000804, 2020P000849, 2015P002215), New York Presbyterian Hospital/Columbia University Medical Center (IRB-AAAT0785, IRB-AAAB2667, IRB-AAAS7370). All patients had confirmed COVID-19 by PCR testing. Consent for autopsy and research was obtained from the healthcare proxy or the next of kin. Massachusetts Institute of Technology (MIT) IRB protocols 1603505962 and 1612793224, and/or the not-involving-human-subjects research protocol ORSP-3635, cover all secondary analyses performed at the Broad Institute of MIT and Harvard. No subject recruitment or ascertainment was performed as part of the Broad protocol. Donor identities and accompanying information were encoded at the relevant hospital site prior to shipping to or sharing with the Broad Institute for sample processing or data analysis. We also included snRNA-seq data from snap-frozen biopsies from 4 healthy neurologically-deceased donor livers suitable for transplantation (G.B., S.A.M), age 40-49 (F), age 40-49 (M), age 40-49 (F), age 20-29 (F) (**Table 1**).

### Sample acquisition

Beth Israel Deaconess Medical Center (BIDMC): Sample collection for BIDMC samples was performed by an interventional radiologist via a 13G coaxial guide with a 14G core biopsy and 20 mm sample length under ultrasound guidance. All biopsies were conducted within 3 hours of confirmed asystole on a gurney in the hospital morgue. All personnel were wearing standard personal protective equipment prior to removing the body from the bag. Multiple biopsies were acquired by tilting the coaxial needle a few degrees in different directions. Core biopsies were separated in two groups: one for formalin fixing and the other flash-frozen in liquid nitrogen and stored at -80 °C until use.

Brigham and Women’s Hospital (BWH): Sample collection for BWH was performed in a negative pressure isolation room with personnel wearing personal protective equipment (powered air-purifying or N95 respirators). Abdominal organs were harvested *en bloc* and the liver was then subsequently dissected, weighted, and photographed. Liver samples were collected from the organ and placed in 25 mL of RPMI-1640 media with 25 mM HEPES and L-glutamine (ThermoFisher Scientific) + 10% heat inactivated FBS (ThermoFisher Scientific) in 50 mL falcon tubes (VWR International Ltd). Tissue samples were transported to Broad in a cooler.

Massachusetts General Hospital (MGH): Sample collection for MGH was performed in a negative pressure isolation room from personnel wearing personal protective equipment (N95 or powered air-purifying respirators). As in BWH, organs were removed *en bloc*, dissected, photographed, and weighed. Liver samples were placed in collection tubes and subsequently in a cooler for transport to the Broad Institute.

New York Presbyterian Hospital: Sample collection was performed as in ^7^. Tissue samples were collected during rapid autopsy within hours from time of death. Tissue samples of ∼1cm^3^ were embedded in Tissue-Tek optimal cutting temperature (OCT) compound (Sakura Finetek USA Inc) and stored at −80⁏°C.

### Tissue processing and single nuclei encapsulation

All samples from all hospitals were snap frozen for the snRNA-seq studies. All sample handling steps were performed on ice. TST and ST buffers were prepared fresh as previously described (Drokhlyansky et al., 2020; Slyper et al., 2020). A 2x stock of salt-Tris solution (ST buffer) containing 292 mM NaCl (ThermoFisher Scientific), 20 mM Tris-HCl pH 7.5 (ThermoFisher Scientific), 2 mM CaCl2 (VWR International Ltd) and 42 mM MgCl2 (Sigma Aldrich) in ultrapure water was made and used to prepare 1xST and TST. TST was then prepared with 1 mL of 2x ST buffer, 6 μL of 10% Tween-20 (Sigma Aldrich), 10 μL of 2% BSA (New England Biolabs), and 984 μL of nuclease-free water 1xST buffer was prepared by diluting 2x ST with ultrapure water (ThermoFisher Scientific) in a ratio of 1:1). 1 mL of PBS-0.02% BSA was also prepared with 990 uL UltraPure 1x PBS ph 7.4 (ThermoFisher Scientific) and 10 uL 2% BSA (New England Biolabs) for sample resuspension and dilution prior to 10x Genomics chip loading. Single frozen biopsy pieces were kept on dry ice until immediately prior to dissociation. With clean forceps, a single frozen biopsy was placed into a gentleMACS C tube on ice (Miltenyi Biotec) containing 2 mL of TST buffer. gentleMACS C tubes were then placed on the gentleMACS Dissociator (Miltenyi Biotec) and tissue was homogenized by running the program “m_heart_02” x 2 until tissue was fully dissociated. A 40 μm filter (CellTreat) was placed on a 50 mL falcon tube (Corning). Homogenized tissue was then transferred to the 40 μm filter and washed with 3 mL of 1xST buffer. Flow-through was transferred to a 15 mL falcon tube (Corning). Samples were then centrifuged at 500g for 5 minutes at 4ºC with brake set to “low”. Sample supernatant was removed and the pellet was resuspended in 100 μL – 200 μl PBS-0.02% BSA. Nuclei were counted and immediately loaded on the 10x Chromium controller (10x Genomics) for single nucleus partitioning into droplets.

### Single nuclear RNA sequencing

For each sample, 8,000-16,500 nuclei were loaded in one channel of a Chromium Chip (10x Genomics). 3’ v3.1 chemistry was used to process all other tissues. cDNA and gene expression libraries were generated according to the manufacturer’s instructions (10x Genomics). cDNA and gene expression library fragment sizes were assessed with a DNA High Sensitivity Bioanalyzer Chip (Agilent). cDNA and gene expression libraries were quantified using the Qubit dsDNA High Sensitivity assay kit (ThermoFisher Scientific). Gene expression libraries were multiplexed and sequenced on an Illumina sequencer.

### SnRNA-seq expression quantification and correction for ambient RNA

The raw sequencing reads were demultiplexed using Cell Ranger mkfastq (10x Genomics). We trimmed the reads from the BIDMC liver samples for polyA tails and the template switching oligo 5’-AAGCAGTGGTATCAACGCAGAGTACATrGrGrG -3’ with cutadapt v.2.7 ^71^. The reads were aligned to generate the count matrix using Cell Ranger count (10x Genomics) on Terra with the cellranger_workflow in Cumulus ^72^. The reads were aligned to a custom-built Human GRCh38 and SARS-CoV-2 (“GRCh38_premrna_and_SARSCoV2’’) RNA reference. The GRCh38 pre-mrna reference captures reads mapping to both exons or introns ^73^. The SARS-CoV-2 viral sequence (FASTA file) and accompanying gene annotation and structure (GTF file) are as previously described ^74^. The GTF file was edited to include only CDS regions, with added regions for the 5’ UTR (“SARSCoV2_5prime”), 3’ UTR (“SARSCoV2_3prime”), and anywhere within the Negative Strand (“SARSCoV2_NegStrand”) of SARS-CoV-2. Trailing A’s at the 3’ end of the virus were excluded from the SARSCoV2 fasta file ^6^. CellBender remove-background ^75^ was run to remove ambient RNA and other technical artifacts from the count matrices. The workflow is available publicly as cellbender/remove-background (snapshot 11) and documented on the CellBender GitHub repository as v0.2.0: https://github.com/broadinstitute/CellBender.

### Filtering of low quality cells and sample integration

We filtered out nuclei with fewer than 400 UMIs, 200 genes, or greater than 20% of UMIs mapped to mitochondrial genes. Furthermore, we discarded samples with less than 100 nuclei. We retained all nuclei that pass the quality metrics described above. Subsequently, snRNA-seq data from individual samples were combined into a single expression matrix and analyzed using Seurat v.3.2.3 ^76–78^. The UMI counts for each nuclei were divided by the total counts for that nuclei, and multiplied by a scale factor of 10,000. Then, values are log-transformed using log1p resulting in log(1+10,000*UMIs/Total UMIs) for each nucleus.

Subsequently, highly variable genes were identified using Seurat’s FindVariableFeatures function. Then, data dimensionality was reduced to the top 15 principal components by PCA using the top 2000 highly variable genes. The lower dimensional embedding was then corrected for technical noise using each sample as a separate batch with Harmony ^79^. Neighbors were computed using the Harmony-corrected embedding. The UMAP and Leiden clusters were computed using the resulting nearest neighbor graph.

### Doublet detection

We used a two-step procedure to identify doublets. First, we identified doublets in each sample with the re-implementation of the Scrublet ^80^ algorithm in Pegasus ^6,72^. Second, we integrated and clustered all samples and identified clusters significantly enriched for doublets. All nuclei in the enriched clusters were flagged as potential doublets.

In brief, we integrated the nuclei that passed the quality control, normalized each nuclei to feature counts per 100K counts (FP100K) and log transformed the expression values (log(FP100k + 1)), selected highly variable genes, computed the first 30 principal components (PCs), corrected the PCs for batch effects using Harmony, and clustered the cells using the Harmony corrected embedding with the Leiden algorithm. Then, we tested if each cluster is significantly enriched for doublets using Fisher extract test controlling at a False Discovery Rate of 5%. Among the significantly enriched clusters, we selected those with more than 60% of nuclei identified as potential doublets and marked all nuclei in these clusters as doublets.

### Clustering

We first derived compartments, high-level clusters, encompassing major cell types. Then, we performed iterative clustering to identify cell types. We used the first 15 PCs corrected by Harmony to compute the nearest neighbor graph. Then we identified the compartments using the Leiden algorithm implemented in the FindClusters function in Seurat. For each compartment, we subsetted the nuclei, selected highly variable genes, computed the first 15 PCs, corrected the PCs for batch effects using Harmony, computed the nearest neighbor graph with the Harmony embedding, and clustered the nuclei using the FindClusters function in Seurat.

### Batch effect correction

Building on approaches that use residuals from a negative binomial generalized linear model (NB-GLM) to normalize single cell data ^81–83^, we fitted a NB-GLM using an efficient implementation of a Gamma-Poisson GLM ^18,84^ with batch as the covariates. We then used the deviance residuals from this model as the expression adjusted for batch effects. For downstream analysis that required counts, we also generated counts corrected for batch by expanding and scaling the model described by ^17^ using a scalable implementation of a Gamma-Poisson GLM ^18^.

### Pathway activity score calculation

A pathway score summarizes the expression of a set of functionally related genes ^85^. A Gene Ontology ^86^ set of 989 GO Biological Process terms was used to create a curated selection of pathways capturing liver parenchymal and non-parenchymal cellular functions and pathways (**Extended Data Table 9**). Building on the methodology described in ^85,87^, we used a rank based approach to define the pathway scores, where the pathway score is the sum of the adjusted ranks of the genes in the pathway annotation scaled by the square root of the number of genes in the pathway. First, the ranks based on the UMI counts are calculated per gene for each nucleus solving ties by selecting the minimum. Then, we scale and center the ranks across each nucleus. In order to account for the effect of rank sparsity for each gene we split the scaled and centered ranks by their sign (positive or negative) and regress out with a linear model the effect of the number of genes detected and the log of the total number of UMIs. Finally, we use the removeBatchEffect function from limma ^88^ to adjust the pathway scores for batch effects. The same approach was used to estimate a score for the curated signatures described by Sánchez-Taltavull *et al*. (proliferating Kupffer cells) ^*37*^, and by Niethamer *et al*. (influenza-injury signature) ^46^.

### Differential expression analysis at cluster level

Differential expression analysis was carried out using limma-trend ^89,90^ to detect cluster gene markers. First, genes expressed in at least 5% of the nuclei of at least one cluster were selected and then UMI counts were normalized using the TMM normalization ^91^ implemented in edgeR v.3.28.1 ^92^. Then, a linear model “∼ Cluster + Batch” was fitted and modeled the mean-variance relationship with the limma-trend method ^89^ and a robust empirical Bayes procedure ^93^. We used contrasts to compare the mean of a given cluster with all others; a gene is considered a cluster marker if the contrast is significant at an FDR < 0.05 and the cluster coefficient is higher than at least 75% of all other clusters. We performed comparisons at two levels: across all compartments (comparing all clusters identified) and within compartments (comparing clusters only from the same high-level cluster). We used limma to fit the same model “∼ Cluster + Batch” on the pathway scores but without the mean-variance trend since the pathway scores are approximately normally distributed. The criteria to select pathway markers were identical to the cluster markers.

### Healthy reference comparison and differential gene expression

We combined the COVID-19 liver nuclei passing QC and were not marked as doublets with the control liver snRNA-seq dataset into a single expression matrix. Similarly to the COVID-19 snRNA-seq analysis, we normalized each nucleus to TP100K and log transformed the expression values (log(TP100k + 1)), selected highly variable genes, computed the first 30 principal components (PCs), corrected the PCs for batch (we considered each sample as a separate batch) using Harmony, and clustered the cells using the Harmony corrected embedding with the Leiden algorithm. We identified 5 high-level compartments in the combined data set. These high-level clusters matched the compartments identified in the COVID-19 liver data. For each high-level cluster the first 15 PCs were corrected for batch effects using Harmony and the nearest neighbor graph was calculated using the Harmony embedding. The nearest neighbor graphs were used to assign each nucleus from the healthy reference to the relevant cluster.

Differential expression analysis was carried out using limma and mean-variance modeling at the observational level (voom) ^89^ after summing nuclei per cluster per sample ^94^, and the linear model “∼ Disease + SVs”, where SVs are surrogate variables estimated with iterative adjusted surrogate variable analysis (IA-SVA) ^95^. The model was fit to estimate the differences between COVID-19 and healthy livers for each cluster. All clusters with at least 3 samples per group with >5 nuclei per sample were included in the analysis.

### Determination of significant changes in cell type proportions

A binomial generalized linear mixed model (GLMM) was utilized to study the differences in cell type abundances between COVID-19 and control livers. Lme4 version 1.1-27.1 was utilized to fit the model ∼ Cluster*Condition + (1|Sample), and emmeans version 1.6.2-1 to compare the odds ratios of COVID-19 vs Control for each cluster (**Extended Data Table 5**).

### Detection of cells with SARS-CoV-2 content above ambient levels

We adapted methods ^75,96,97^ previously described in ^6^ to designate a single nucleus as SARS-CoV-2 RNA+ or SARS-CoV-2 RNA-. A permutation test was utilized to determine the probability that the nucleus contained a higher SARS-Cov-2 UMI content than expected by ambient contamination, while taking into account the fractional abundance of SARS-Cov-2 aligning UMIs, the abundance of SARS-Cov-2 aligning UMIs in the ambient pool, and the estimated ambient contamination of the single nucleus.

The fractional abundance of SARS-Cov-2 aligning UMIs per nucleus was defined as the number of UMIs assigned to all viral genomic features divided by the total number of UMIs aligning to either the SARS-Cov-2 or GRCh38 reference. The abundance of SARS-Cov-2 UMIs in the ambient pool was defined as the sum of all SARS-Cov-2 UMIs in the pre-CellBender output within discarded nuclei flagged as “empty” or “low quality”. Hence, the ambient fractional abundance was determined for each sample independently. The discarded nuclei were resampled to generate the null distribution of the SARS-CoV-2 fractional abundance, which was utilized to extract empirical p-values for the observed fractional abundance of each nucleus. The empirical p-values were adjusted for multiple comparisons using false discovery rate. Nuclei with at least 2 SARS-Cov-2 UMIs and an FDR < 0.05 were assigned as “SARS-CoV-2 RNA+”; “SARS-Cov-2 Ambient” if having SARS-CoV-2 UMIs but were not significantly higher than the ambient pool; and “SARS-CoV-2 RNA-” if no SARS-Cov-2 UMIs were detected.

### Differential expression analysis between SARS-Cov-2 RNA+ and SARS-Cov-2 RNA-nuclei

In order to test the genes and pathways associated with the presence of SARS-Cov-2 RNA, we used the following approach to account for the biases due to differences in number of nuclei, quality and sample-to-sample variability. First, we only considered cell types with at least 10 SARS-Cov-2 RNA+ nuclei (above ambient levels) and within a given cell type we only considered samples with at least 2 SARS-Cov-2 RNA+ nuclei. Then we subsampled the SARS-Cov-2 RNA-nuclei to match the complexity distributions. The nuclei were partitioned into 5 bins based on complexity, log10(Number of genes/nuclei), and the SARS-Cov-2 RNA-nuclei were subsampled to match the distribution of the SARS-Cov-2 RNA+ nuclei ^8^. We resampled the pool of SARS-Cov-2 RNA-nuclei to generate the null distribution for the mean expression and the pathway scores in order to estimate an empirical p-value for the mean expression in the SAR-Cov-2 RNA+ nuclei. Mean expression was calculated by normalizing the UMI counts using the trimmed mean of M-values (TMM) normalization ^91^ and adjusted for batch effects using limma’s removeBatchEffect function. Pathway scores were estimated for the selected nuclei and then adjusted for batch effects using limma’s removeBatchEffect function. P-values were adjusted for multiple comparisons using FDR.

### Viral enrichment analysis

A viral enrichment score per cluster was calculated as previously^6,98^. The enrichment score for a given cluster C is defined as: EnrichmentI = log((Observed(Vcells in C) + ε) / (Expected(Vcells in C) + ε)) = log((Vcells in C) + ε) / ((Vcells in total * X_c) + ε) where Vcells are the SARS-Cov-2 RNA+ nuclei, X_c is the proportion of the total number of nuclei in cluster C out of the total number of nuclei in its corresponding compartment, and ε = 0.0001. We only considered samples with at least 5 SARS-Cov-2 RNA+ nuclei. We derived the null distribution of each enrichment score by permuting the data and assigning the same number of SARS-Cov-2 RNA+ labels to nuclei, such that the overall proportion of SARS-Cov-2 RNA+ nuclei was fixed, computing the cluster enrichment score and estimating the empirical p-value as the fraction of the permutations that showed a similar or higher enrichment score compared to the observed enrichment score. Then, we adjusted the empirical p-values for multiple comparisons using FDR.

### Trajectory interference and cell-cell communication analysis

Single-cell pseudotime trajectory was constructed using Slingshot (version 2.0.0) based on the Harmony embedding matrix. The embedding matrix was re-computed for the Hepatocyte and Biliary Epithelial cells, excluding the BEC3 doublet cluster, while the first 20 dimensions were utilized for the subsequent analysis. Lineages were determined and mapped to the UMAP embedding matrix using the relevant Slingshot protocol ^99^. Cell-cell communication among the distinct cell populations was defined using the CellChat R package ^100^. The average gene expression per cell group was calculated by applying a threshold of 20% and using the batch corrected counts. Significant ligand-receptor interactions and pathways were retained by applying a 0.05 *P value* cutoff.

### Digital Spatial Profiling

Liver tissue sections of 5 μm were prepared from formalin-fixed paraffin-embedded blocks. Tissue integrity was confirmed on slides stained with hematoxylin and eosin (H&E). Slides were stored in vacuum at 4 °C to preserve RNA integrity. To prepare the slides for digital spatial profiling (DSP), slides were stained against Pan-Cytokeratin, CD68, CD45, and DNA. A Whole Transcriptome Atlas (WTA) probe library (Nanostring) was applied on each slide according to the manufacturer instructions. Four categories of area of interest (ROI) for transcriptome profiling were manually selected under a fluorescence-microscope: portal area, and lobular zones 1-3.

Specifically, autopsy FFPE tissues from COVID-19 infected patients were processed following the GeoMx DSP slide prep user manual (MAN-10087-04). Autopsy slides were baked in an oven at 65°C for an hour and then they were processed on a Leica Bond RX automation platform with a protocol including three major steps: 1) slide baking, 2) antigen Retrieval 20min at 100°C, 3) 1.0ug/ml Proteinase K treatment for 15min. Subsequently, the slide was incubated with the RNA probe mix (WTA and COVID-19 spike-in panel, **Extended Data Table 2**). After overnight incubation, slides were washed with buffer and stained with CD68-594 (Novus Bio, NBP2-34736AF647), CD45-647 (Novus Bio, NBP2-34527AF647), PanCK-488 (eBioscience, 53-9003-82) and Syto83 (ThermoFisher, S11364) for 1 hour, and loaded on the NanoString GeoMx DSP to scan 20X fluorescent images. Regions of interest (ROIs) were placed by an expert panel comprising hepatologists, pathologists, and technology specialists. Portal, periportal, Zone 1, 2, and 3 regions were prioritized. Following ROI selection, oligos were then UV-cleaved and collected into 96-well plates. Oligos from each ROI were uniquely indexed using Illumina’s i5 x i7 dual-indexing system. 4 μL of a GeoMx DSP sample was used in the PCR reaction. PCR reactions were purified with two rounds of AMPure XP beads (Beckman Coulter) at 1.2x bead-to-sample ratio. Libraries were paired-end sequenced (2×75) on a NovaSeq 6000 sequencer. Serial sections were subjected also to RNA *in situ* hybridization assay using the RNAScope platform (ACD) and by following the standard vendor protocol.

### NanoString GeoMx DSP data preprocessing

Sequencing reads were compiled into FASTQ files corresponding to each region of interest (ROI) using bcl2fastq. FASTQ files were demultiplexed and converted to Digital Count Conversion (DCC) files with NanoString’s GeoMx NGS DnD Pipeline. The resulting DCC files were converted to an expression count matrix. Raw probe data for 18,372 endogenous genes, with 18,346 genes having one probe per gene and 26 SARS-CoV-2 related genes having 5 probes per gene, as well as 105 global negative probes and 8 SARS-CoV-2 negative probes were generated for 71 ROIs, spanning the portal region, all 3 lobular zones and CD45 regions from 4 patients. The probe counts were normalized using the TMM normalization method implemented in edgeR v.3.28.1. In order to account for unwanted variation, we estimated surrogate variables (SVs) using Iteratively Adaptive Surrogate Variable Analysis (IA-SVA) ^95^ specifying the model “∼ Region + Donor”. The expression values were subsequently adjusted with limma’s removeBatchEffect function with Donor as batch and the SVs as covariates.

### Integration of snRNA-seq and DSP data

The data from the nanoString DSP assay were utilized to infer the location of the clusters identified in the snRNA-seq data using the caret (6.0.90) and RandomForest (4.6.14) packages in R 4.0.1. A random forest classifier was trained to predict whether a sample was located in the lobule or in the portal area using pathway activity scores (PAS) as features. The top 200 differentially activated pathways between portal and lobule (100 most upregulated and 100 most downregulated) identified in the nanoString GeoMx DSP data were incorporated as features in the classifier. PAS were estimated, corrected for batch effects, scaled and centered after summing the nuclei per sample in each cluster. For training, clusters which could be assigned to the lobular or portal area after expert curation were utilized, such as hepatocyte clusters in the lobule and cholangiocytes (BECs) in the portal area. Identified clusters were pseudobulked to reduce noise, and class imbalance was resolved using SMOTE ^101^, owing to the fact that lobular hepatocyte cells significantly outnumbered portal cells. The samples were split into an 80% training set (224 lobular and 168 portal) and a 20% testing set (30 lobular and 13 portal). Optimal training parameters were identified using 5-fold cross validation on the training set through the caret package, resulting in an area under the curve (AUC) of 0.984. Then, the classifier was applied to the remaining clusters. Utilizing SMOTE to address class imbalance, similar results were obtained at the single cell level (Training and CV set: 6,944 Lobular and 5,208 Portal cells after upsampling, Testing Set: 10,778 Lobular and 434 Portal, resulting in an AUC of 0.998).

### NanoString GeoMx DSP pathway activity scores

We also used a rank based approach to estimate pathway scores. First, we established ranks based on the raw probe counts for each ROI. Then, the ranks were centered and scaled (per ROI). We estimated the pathway score as the sum of the scaled and centered ranks of the genes in the pathway annotation scaled by the square root of the number of genes in the pathway. We accounted for unwanted technical variation in the pathways scores by estimating SVs using the IA-SVA method with the model “∼ Region + Donor + log(Nuclei Counts) + log(ROI size)”. Then, we adjusted the pathway scores with limma’s removeBatchEffect function with Donor as batch, the SVs, log(Nuclei Counts), and log(ROI size) as covariates.

### NanoString GeoMx DSP viral scores

A SARS-CoV-2 viral score was calculated for the GeoMx DSP WTA ROIs using the extended SARS-COV-2 probe set. In particular, the probes for the S and ORF1ab SARS-CoV-2 genes were utilized. First, the ranks per ROI were calculated based on the raw counts for both the target and negative probes in the SARS-COV-2 probe set, and subsequently centered and scaled. Following a similar approach to the pathway activity scores, the viral score was calculated as the sum of the scaled and centered ranks for the S and ORF1ab probes multiplied by the square root of 2 (the number of genes in the set). Then, the negative and target probe labels were permuted 10,000 times and the viral score was calculated for each permutation to estimate the mean and standard deviation of the viral score. Using these estimates, the observed viral score in each ROI was centered and scaled. Limma’s removeBatchEffect function with the model “log(Nuclei counts) + log(ROI size)” as covariates was utilized to account for ROI size and nuclei counts within the ROI. Finally, the adjusted viral scores were fit to the linear model “∼ 0 + Donor” using limma to compare the viral scores between donors. For each donor, a contrast was fit to compare the mean adjusted viral score with the mean of the other donors. For example, the contrast for donor L1 is “Donor L1 - (Donor L2 + Donor L3 + Donor L4)/3”.

### NanoString GeoMx DSP differential expression analysis

Limma-trend was utilized to perform differential expression analysis with the GeoMx DSP data. First, batch-corrected expression was fit into the model “∼ Region” with the limma-trend method and a robust empirical Bayes procedure. Contrasts were utilized to compare the mean of a region against all others, with a gene considered as a region-specific marker if the contrast was significant at an FDR of 0.05 and the region coefficient higher than all other regions. Limma was also used to fit the same model “∼ Region” on the pathway scores but without the mean-variance trend since the pathway scores are approximately normally distributed. The criteria to select pathway markers were the same as for genes.

For the rotation/scale normalized zonation gradient, ROIs were grouped by lobule and the distance to the zone 1 ROI was calculated per ROI, per lobule. Distances were normalized to be in the [0,1] range. Using the normalized distances, the model “∼ Normalized Distance” was fit with the batch corrected values, the limma-trend method, and a robust empirical Bayes procedure. We used the coefficient for the normalized distance to identify genes that have an increasing and decreasing pattern across the zonation gradient. For the pathway scores, the same model was fit without the mean-variance trend.

#### Quantitative RT-PCR against SARS-CoV-2

Total RNA was extracted from liver tissue samples using a QIAcube HT (Qiagen) and RNeasy 96 QIAcube HT Kit (Qiagen). RNA was reverse transcribed into cDNA with superscript VILO (Invitrogen). SARS-CoV-2 N (nucleocapsid) gene was cloned into a pcDNA3.1 expression plasmid and transcribed using an AmpliCap-Max T7 High Yield Message Maker Kit (Cellscript) to be utilized as a standard. qPCR was performed in duplicates using a QuantStudio 6 Flex Real-Time PCR System (Applied Biosystems). Viral load was calculated as RNA copies per microgram of total RNA, with a quantitative assay sensitivity of 50 copies. Primers utilized for SARS CoV-2 N genes were: 2019-nCoV_N1-Forward : 5’-GACCCCAAAATCAGCGAAAT-3’, 2019-nCoV_N1-Reverse: 5’-TCTGGTTACTGCCAGTTGAATCTG-3’, and probe: 2019-nCoV_N1-Probe: 5’-FAM-ACCCCGCATTACGTTTGGTGGACC-BHQ1-3’

#### Subgenomic mRNA assay

SARS-CoV-2 E gene subgenomic mRNA (sgmRNA) was assessed by RT-PCR as in Wölfel *et al*. ^102^. A Taqman custom gene expression assay (ThermoFisher Scientific) was utilized to target the E gene sgmRNA ^102^. Standard curves were used to calculate sgmRNA in copies per microgram of total RNA with an assay sensitivity of 50 copies.

### RNAScope

RNA in situ hybridization (ISH) was performed with the RNAScope Multiplex Fluorescent Kit (ACDBio, Newark, CA). All three probes (Hs-TMPRSS2, Hs-ACE2-C2, V-nCoV2019-S-C3) were designed by ACDBio to ensure target specificity. FFPE liver biopsy sections at 5 μm were first de-paraffinized using xylene and ethanol, and incubated in the pretreatment buffer with protease and incubated in a HybEZ oven (ACDBio). The staining of mRNA was achieved by hybridization with the target probes over the pretreated liver tissue, followed by sequential treatment of amplification reagents provided in the RNAScope kit. Each section was dehydrated before being mounted with Pertex (ACDBio). A probe against a housekeeping gene PPIB was used as a positive control (ACDBio).

### Histology, immunohistochemistry, and special tissue staining

Connective tissue stain (Sirius red) and immunohistochemistry (IHC) were performed using formalin-fixed, paraffin-embedded liver biopsy of four COVID-19 patients. For Sirius red staining, liver sections were dewaxed, rehydrated and stained for 2 minutes with hematoxylin, then 30 minutes with a picrosirius red solution (ab246832, Abcam). For IHC staining, antigen retrieval of dewaxed and rehydrated paraffin-embedded liver sections was performed using sodium citrate pH=6 for α-SMA and pepsin for CK19, respectively, followed by blocking with 10% goat serum for 30 minutes, and incubation with anti-α-SMA (Cell Signaling Technology, 19245, 1:400) and anti-CK19 (Sigma-Aldrich, MAB3238, 1:100) primary antibody overnight at 4 °C. After incubation with biotinylated secondary antibody for 1.5 hours, detection was performed with the Vectastatin Elite ABC-HRP kit (Vector Laboratories, SP-6100) with the DAB Peroxidase Substrate kit (Vector Laboratories, SK-4100), followed by counterstaining with haematoxylin.

## Author Contributions

Z.G.J., Y.P., A.R., A.K.S., B.I., and I.S.V. conceived and led the study. W.H., G.S., J.H., O.R.R., L.T., A-C.V, O.A., M.B., D.H, C.P., S.R., I.H.S, N.I., G.D.B, S.A.M., R.F.P, Z.G.J., Y.P., A.R., A.K.S., B.I., and I.S.V. supervised the study. Y.P.J., D.K., N.K., P.N., T.H., T.S.A., C.G.J.Z., J.R., A.S., and S.J.F performed the data analyses. J.C.M., A.L.E., D.P., D.B., P.H., L.P., A.D.A., J.B., H.H., M.V., Z.K., C.J., T.M.D, D.P., Z.B.A., V.M.T, J.G., A.S., S.Z, M.S and C.G.J.Z. contributed in sample preparation and the performance of the spatial/single cell experimental studies. J.H., O.B., I.H.S., R.F.P. contributed to the sample procurement and preparation. Z.G.J., Y.P., I.S.V., S.N., A.M., R.C., N.I., N.K., D.K., L.P., J.H., and Y.P.J contributed to the cluster/ROI/slide annotation. J.B., S.W., E.M., J.R., T.H., L.P., Y.P., and Z.G.J. contributed to the performance of the spatial assays. Y.P.J., D.K., N.K. guided analysis, and contributed methods.

## Acknowledgements

We are deeply grateful to all donors and their families. We thank all members of the Departments of Pathology in the relevant medical centers (Beth Israel Deaconess Medical Center, Brigham and Women’s Hospital, Massachusetts General Hospital, Columbia University Irving Medical Center) who led the procurement of autopsy tissues used in this work. NanoString for early access to the WTA assay and technical support as well as 10x Genomics and Illumina for support during the data generation process. Portions of this research were conducted on the Ithaca High Performance Computing system, Department of Pathology, BIDMC, and the O2 High Performance Compute Cluster at Harvard Medical School. The project has been funded in part with funds from the Manton Foundation, Klarman Family Foundation, HHMI, the Chan Zuckerberg Initiative, and the Human Tumor Atlas Network trans-network projects SARDANA (Shared Repositories, Data, Analysis and Access). A.R. was an Investigator of the Howard Hughes Medical Institute. A.K.S. is supported by US Food and Drug Administration grant HHSF223201810172C, Sloan Fellowship in Chemistry, the Ragon Institute, the Bill and Melinda Gates Foundation (OPP1202327, INV-027498). I.S.V. is supported by the National Institute of Health (NIH) National Cancer Institute grants R01 CA258776, R01 HL129506-07, National Blood Clot Alliance, and Department of Defense PR200524P1. The BIDMC Spatial Technologies Unit was supported by a Research Infrastructure Program Award, Massachusetts Life Sciences Center. B.I. is supported by NIH NCI grants K08CA222663, R37CA258829, R21CA263381, U54CA225088, a FastGrant, the Burroughs Wellcome Fund Career Award for Medical Scientists, and the Louis V. Gerstner, Jr. Scholars Program. A.K.S. is supported by US Food and Drug Administration grant HHSF223201810172C, Sloan Fellowship in Chemistry, the Ragon Institute, the Bill and Melinda Gates Foundation (OPP1202327, INV-027498). This research was funded in part through the NIH Support Grant S10RR027050 for flow cytometry analysis and the NIH/NCI Cancer Center Support Grant P30CA013696 at Columbia University Genetically Modified Mouse Model Shared Resource, Molecular Pathology Shared Resource and its Tissue Bank.

## Conflict of Interest

A.K.S. reports compensation for consulting and/or SAB membership from Merck, Honeycomb Biotechnologies, Cellarity, Repertoire Immune Medicines, Ochre Bio, Third Rock Ventures, Hovione, Relation Therapeutics, FL82, Senda Biosciences, Empress Therapeutics, IntrECate Biotherapeutics, and Dahlia Biosciences unrelated to this work. A.R. is a founder and equity holder of Celsius Therapeutics, an equity holder in Immunitas Therapeutics and until August 31, 2020 was an SAB member of Syros Pharmaceuticals, Neogene Therapeutics, Asimov and ThermoFisher Scientific. From August 1, 2020, A.R. is an employee of Genentech, a member of the Roche Group, with equity in Roche. I.S.V. consults for Guidepoint Global, Mosaic, and NextRNA. G.S. consults in Alnylam Pharmaceuticals, Merck, Generon, Glympse Bio, Inc., Mayday Foundation, Novartis Pharmaceuticals, Quest Diagnostics, Surrozen, Terra Firma, Zomagen Bioscience, Pandion Therapeutics, Inc., Durect Corporation; royalty from UpToDate Inc. and Editor service for Hepatology Communications. Y.P. receives grant support from Enanta Pharmaceuticals, CymaBay Therapeutics, Morphic Therapeutic; consulting and/or SAB in Ambys Medicines, Morphic Therapeutics, Enveda Therapeutics, BridgeBio Pharma, as well as being an Editor of American Journal of Physiology-Gastrointestinal and Liver Physiology. Z.G.J. receives grant support from Gilead Science, Pfizer and compensation for consulting from Olix Pharmaceuticals. L.P, T.H., S.W, J.B, E.M, and J.R. are employees and/or stockholders of NanoString Technologies. B.I. has received honoraria from consulting with Merck, Janssen Pharmaceuticals, AstraZeneca, and Volastra Therapeutics. A.R. and O.R.R. are inventors on multiple patents from the Broad Institute related to single cell and spatial genomics.

## Tables

**Table 1.** COVID-19 and control cohort overview. (PMI: post-mortem interval).

## Extended Data Tables

**Extended Data Table 1:** Liver serum markers for COVID-19 and control liver samples.

**Extended Data Table 2:** NanoString GeoMx Digital Spatial Profiling (DSP) Whole Transcriptome Atlas (WTA) SARS-CoV-2 additional probe set.

**Extended Data Table 3:** Significant genes and pathways following the zonation gradient in the GeoMx DSP WTA data.

**Extended Data Table 4:** Differentially expressed genes and pathways for each region of interest in the GeoMx DSP WTA data.

**Extended Data Table 5:** Differential abundance results comparing COVID with Control livers using a Binomial generalized linear mixed model (GLMM).

**Extended Data Table 6:** Summary of viral loads using RT-PCR and subgenomic mRNA assay in donors L1-5 who expired due to COVID-19 (LOQ: limit of quantification).

**Extended Data Table 7:** Association between clinical covariates and donor viral enrichment score. The association tests (Spearman correlation, Kendall’s tau and Wilcoxon rank sum test) were conducted for all samples from all medical centers (All.statistics, All.pvalue) as well as for the samples from medical center A separately (CentrerA.statistic, CenterA.pvalue).

**Extended Data Table 8**: Summary of liver histopathology findings for samples L1 to L4. H&E staining, CK19, and α-SMA IHC, as well as connective tissue staining (picrosirius red) were performed in four consecutive core biopsies samples and evaluated by an expert clinical liver pathologist (I.N.).

**Extended Data Table 9:** Curated pathway annotations and signatures used to estimate pathway activity scores.

## Extended Data Figures

**Extended Data Figure 1:**
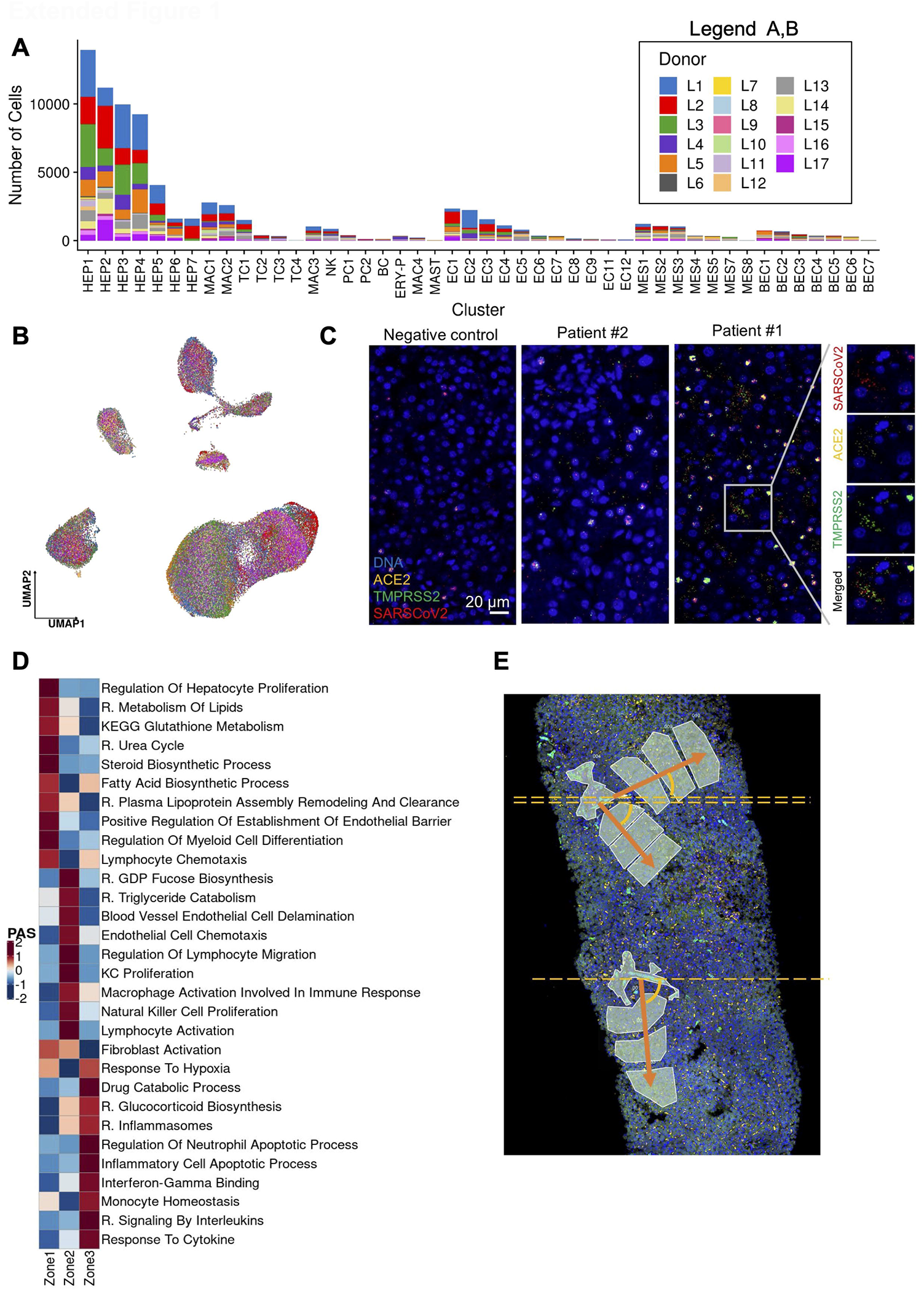
(A) Number of cells per donor for each cluster. Donors are marked with distinct colors. (B) Uniform manifold approximation and projection (UMAP) of all COVID-19 patient liver cells colored per donor. (C) Liver biopsy tissue (5 μm section) from Donors L1, 2 and a control sample processed with the RNAScope Fluorescent Multiplex Assay (Biotechne). Green: TMPRSS2 (Hs-TMPRSS2); Yellow: ACE2 (Hs-ACE2-C2); Red: SARS-CoV-2 (V-nCoV2019-S-C3); Blue: DAPI. Magnified panels (right) show the single channel staining of each probe in Donor L1. Scale bar represents 20 μm. (D) Heatmap of pathways exhibiting a zonated activity gradient in the DSP WTA data. The zonated pathways were determined by regressing the normalized distance to the zone 1 ROI with the pathway activity score. Color denotes the average pathway activity score of all regions of interest (ROIs) collected for each lobular zone following normalization for batch. Displayed pathways are derived from GO, Biocarta (B), and Reactome (R). (E) Zonation distance diagram depicting the rotation/scale invariant modeling applied for the calculation of the pathway activity score gradient. The ROIs were grouped by lobule and the distance to the corresponding zone 1 ROI was normalized to the (0,1) range. The normalized distance accounts for differences in scale and orientation.

**Extended Data Figure 2:**
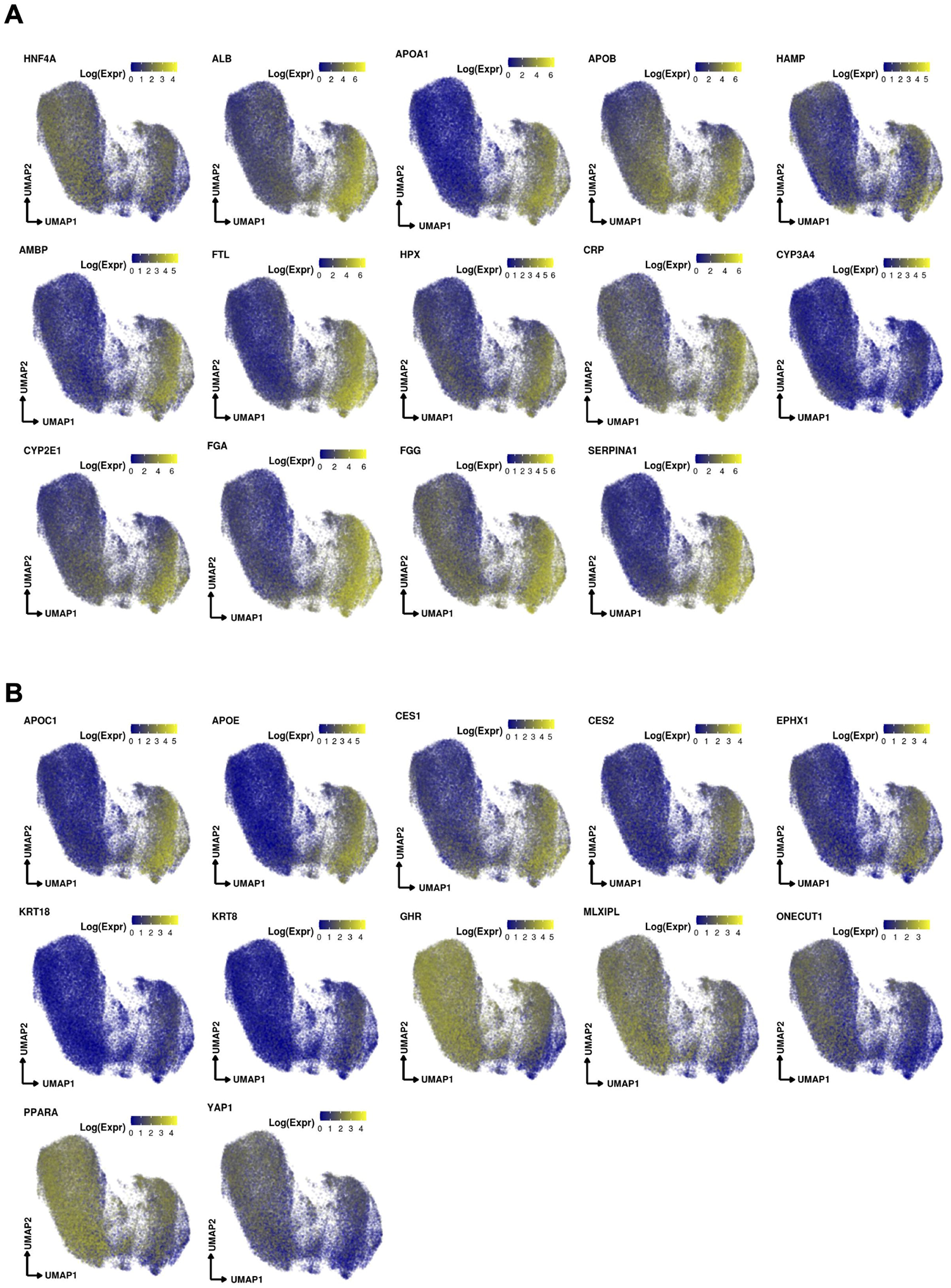
(A) Uniform manifold approximation and projection (UMAP) depicting gene marker expression in the Hepatocytes compartment. (B) UMAP of markers with higher expression in the left or right portions of the Hepatocyte compartment, denoting a potential division of labor.

**Extended Data Figure 3:**
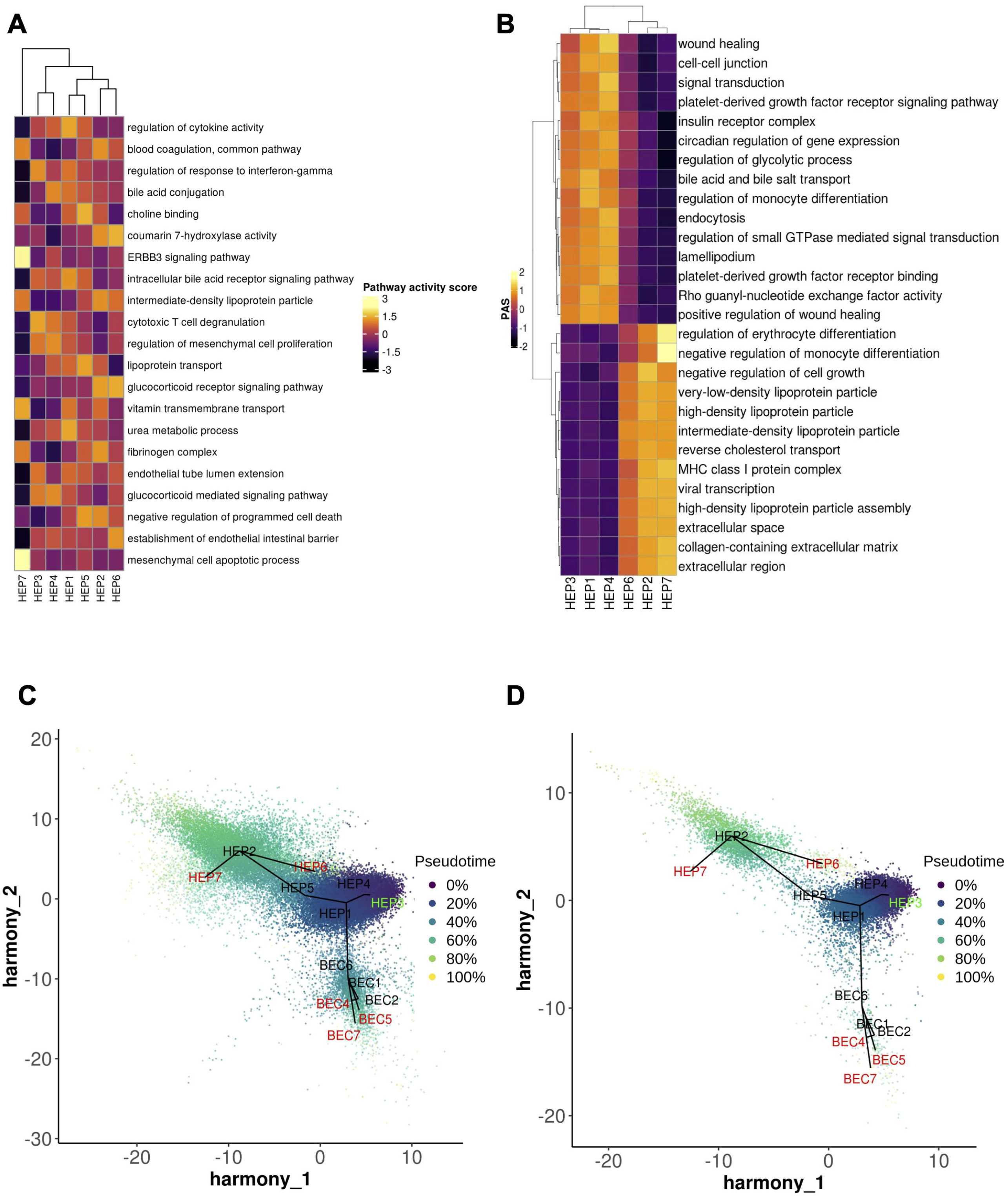
(A-B) Heatmaps capturing highly active pathways based on pathway activity scores in (A) Hepatocytes and (B) Right versus Left Hepatocyte compartment cellular populations. (C-D) Pseudotime analysis using Slingshot. Cells are colored based on pseudotime values and are projected on the first 2 primary harmony embeddings across 5 lineages of Hepatocyte and Biliary epithelial cells for (C) COVID-19 and (D) healthy liver samples. The initiating and terminal lineage nodes are represented with green and red, respectively.

**Extended Data Figure 4:**
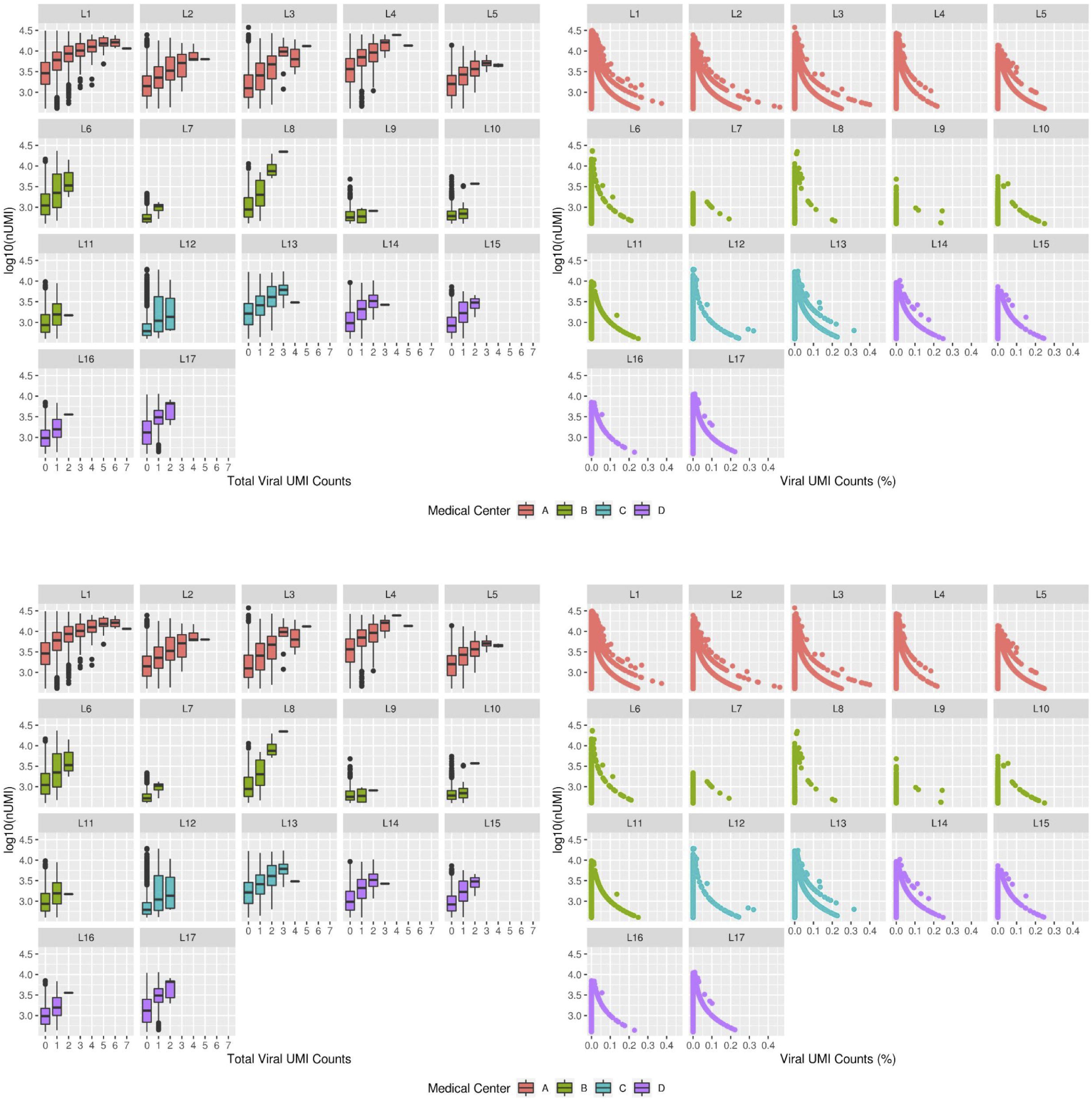
Viral UMI counts as a function of the number of genes (nGenes) or total UMIs (nUMI) detected in the snRNA-seq data across all donors. Left: Boxplots per viral UMI count depicting the number of detected genes (top) or total UMIs (bottom) on a log10 scale. Right: Scatterplots of % viral UMI counts per cell vs the number of detected genes (top) or total UMIs (bottom) on a log10 scale.

**Extended Data Figure 5:**
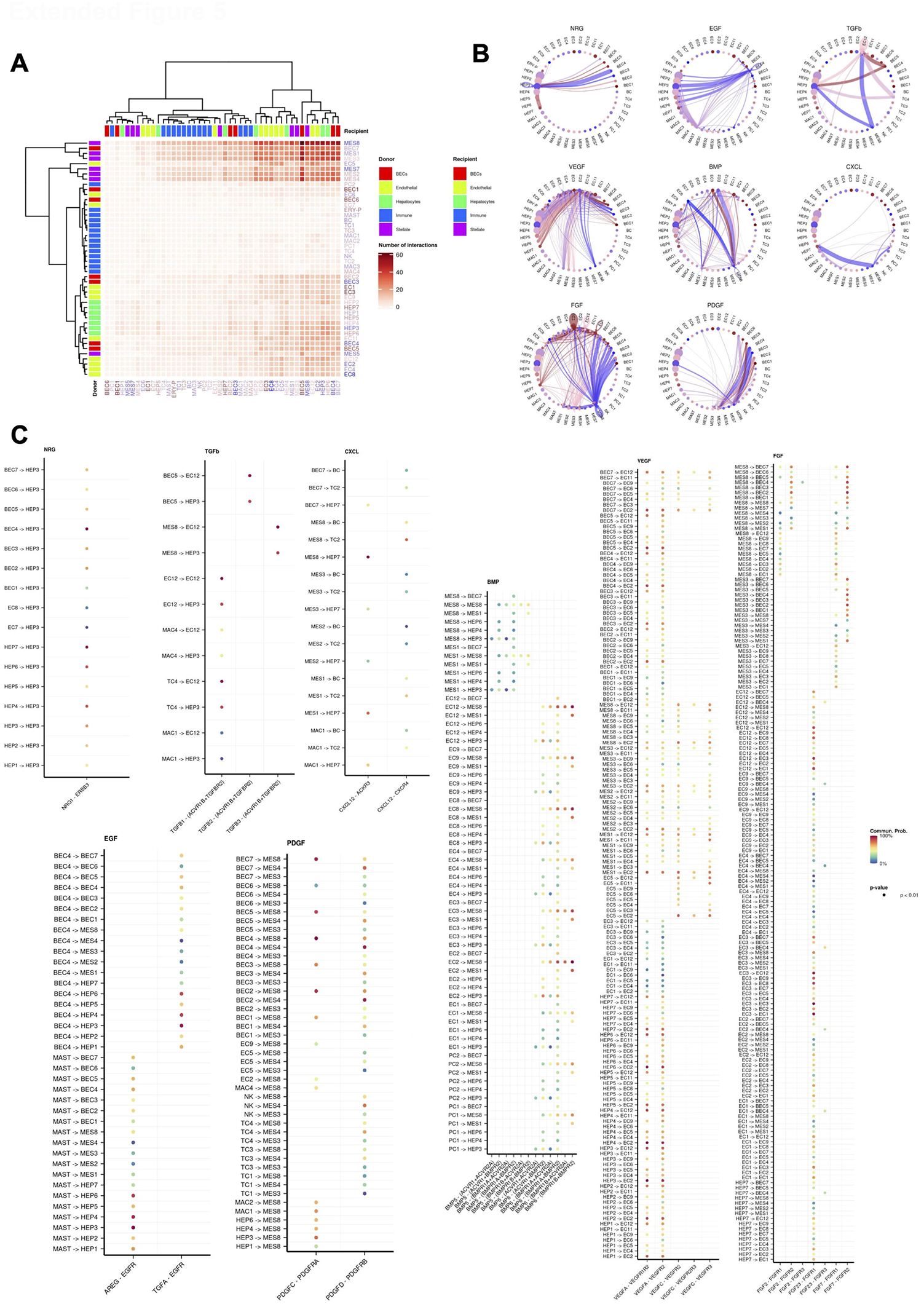
(A) Heatmap portraying cell-cell communication between the detected cell clusters. The color gradient indicates the number of interactions identified between any two cell groups. Recipient/Donor cell-type color is portrayed in a blue (healthy) to red (COVID-19) gradient, concordantly with the cell composition fold-change differences between healthy and COVID-19 liver samples. (B) Circle plots portraying the aggregated cell-cell communication network in NRG, EGF, TGFb, VEGF, BMP, CXCL, FGF, and PDGF pathways. A thicker edge line indicates a stronger signal, while circle sizes are proportional to the number of cells in each cellular compartment. Donor edge-line and circle color are portrayed in a blue (healthy) to red (COVID-19) gradient, concordantly with the cell composition fold-change differences between healthy and COVID-19 liver samples. (C) Dot plots depicting the relative communication probability of each ligand-receptor (x-axis) in any two significantly interacting cellular compartments (y-axis) (*P-value* < 0.01) for NRG, EGF, TGFb, CXCL, BMP, VEGF, FGF, and PDGF pathways. Lowest to highest relative communication probability is portrayed with a blue to red color gradient.

**Extended Data Figure 6:**
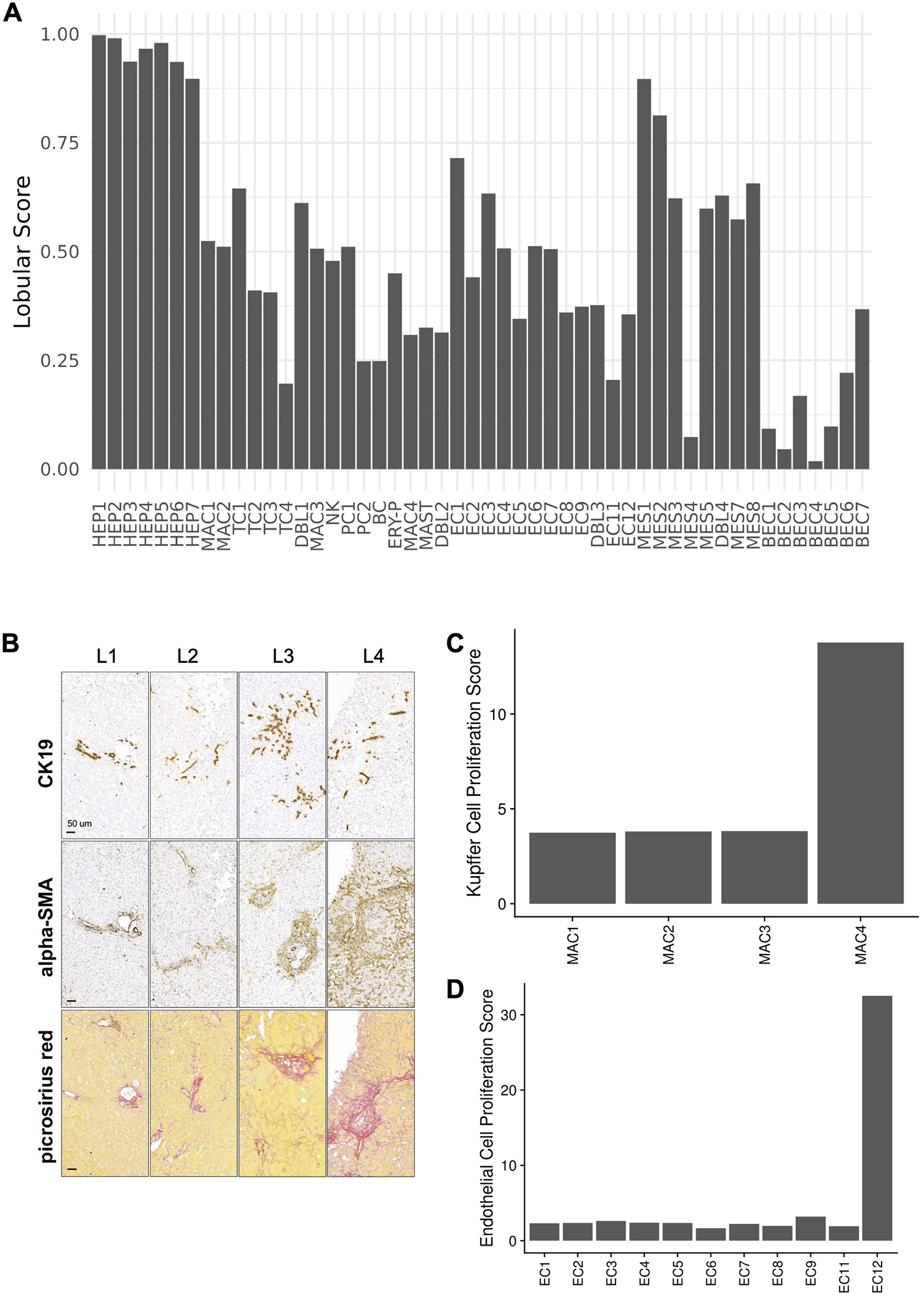
(A) Probability of each snRNA-seq cluster being localized in the hepatic lobular region based on the DSP WTA data. (B) Representative images of serial sections from four consecutive liver core biopsies samples (BIDMC cohort, donors L1 to L4 as indicated on each column) stained for the ductular/cholangiocyte cell marker CK19, HSC activation maker α-SMA, and connective tissue (picrosirius red), as indicated. All images were acquired at the same magnification (scale bar is 50um). (C) Pathway activity score for the Kupffer cell proliferation signature described by ^37^ for the macrophage clusters of the immune compartment. MAC4 was characterized as Replicating Kupffer Cells. (D) Pathway activity score for the endothelial cell proliferation signature described by ^46^ for the Endothelial cell clusters. EC12 was annotated as Replicating Endothelial Cells.

