## Supplementary Note for "A single-nucleus and spatial transcriptomic atlas of the COVID-19 liver reveals topological, functional, and regenerative organ disruption in patients"

**Cell Population Dictionary**

**Hepatocytes (HEP)**

Hepatocyte nuclei accounted for 63.8% of all nuclei, were split into 7 distinct subclusters, and spanned a continuum between two dichotomous ends: primary essential liver functions, such as production of blood proteins, and cell differentiation and replenishment alongwith response to stress.

**HEP1:** 27% of hepatocytes. Interzonal hepatocytes. Low metabolic activity and high activity in pathways relating to cellular differentiation, growth and wound healing (HNF4A, HNF4B, YAP/TAZ, PPRA/B/G, STAT3 and GHR. Gene expression indicates this cluster is composed of hepatocytes with bile related activity, as they are ABCC2h and ONECUT1 high. Potentially a key intermediate between HPC and both hepatocytes and cholangiocytes. Similar proportion between healthy and COVID-19 liver samples.

**HEP2:** 21.7% of hepatocytes. Mature, metabolically active hepatocytes. Increased expression of genes encoding circulating blood proteins, consistent with this subcluster containing mature, differentiated hepatocytes. Also enriched in expression of genes related to APO, iron metabolism and alcohol metabolism. Cell-Cell Communication Analysis unveiled the importance of the LIGHT and CXCL pathways in the CCI of HEP2 and HEP5. Significantly enriched population in COVID-19 liver samples.

**HEP3:** 19.3% of hepatocytes. Least differentiated cells, with high expression of growth factor genes, related to the WNT and NOTCH pathways, suggesting that this subcluster contains cells with activated growth programs. Trajectory analysis indicated a pseudotime path between HEP3 to HEP2, with HEP1, 4 and 5 as intermediates. Significantly depleted population in COVID-19 compared to healthy samples. Significantly enriched in viral reads.

**HEP4:** 17.9% of hepatocytes. Hepatocytes indicating stress response. Low HNF4A, APOB and high SCARB1 and ARG1 expression, as well as stress related pathways, like heat shock, NF-κβ and hypoxia., consistent with a phenotype identified with bulk proteomics on severe COVID-19 patient liver samples. Furthermore, HEP4 nuclei had high expression of collagen modifying enzymes and VEGF-A, indicating potential regulation of hepatocyte-endothelial interactions. Enriched in Zone3 according to GeoMX DSP. Significantly enriched population in COVID-19 liver samples compared to healthy samples. Significantly enriched in viral reads.

**HEP5:** 7.9% of hepatocytes. Substantial CD45 expression compared to other hepatocyte clusters, reported to be a characteristic of hepatocyte progenitors without other standard progenitor characteristics in high abundance. Significantly enriched population in COVID-19 liver samples compared to healthy samples. Increased expression of MT genes.

**HEP6:** 3.1% of hepatocytes. High MT hepatocytes. Similar gene expression profile to HEP2. Distinct footprint related to apoptosis and senescence. Trajectory analysis revealed a path from HEP2 to HEP6. Significantly enriched population in COVID-19 liver samples, as compared to healthy. Significantly enriched in viral reads.

**HEP7:** 3.1% of hepatocytes. Acute response hepatocytes. Similar gene expression profile to HEP2, with marked increase in acute phase protein expression (CRP, C3, C4α, SAA1, and FTH1). CCI analysis revealed that cells in this subcluster interact with cells in the mesenchymal compartment (MES8), as well as having increased expression of CXCL12 pathway ligands and receptors. Related to HEP2 in pseudotime. This subcluster was absent from healthy samples. Significantly enriched in viral reads.

**Biliary Epithelial Cells (BEC)**

Biliary epithelial cell nuclei accounted for 3.6% of all nuclei and were split into 7 main representing both differentiated cholangiocytes, hepatocyte progenitor cells and a profibrogenic reactive cholangiocyte subject, along with a possible artifact subcluster. The subclusters expressed the lineage markers CFTR, KRT7 and KRT19.

**BEC1:** 25.1% of Biliary Epithelial Cells. Small mature cholangiocytes. Consistent with fully differentiated small size cholangiocytes that line small caliber bile ducts, expressing SCTR, BCL2 and primary cilia genes (BICC1, PKHD1, DCDC2, CTNND2, PKD2, but not CYP2E1). Significantly enriched population in COVID-19 liver samples.

**BEC2:** 23.4% of Biliary Epithelial Cells. Small mature cholangiocytes. A second subcluster of fully differentiated small size cholangiocytes, but had higher PDGFD, ZNK19, PAK3, ONECUT1, and CD133 compared to BEC1. Significantly enriched population in COVID-19 liver samples.

**BEC3:** 15.6% of Biliary Epithelial Cells. Potentially doublets High MT gene expression, as well as hepatocyte specific markers, like CPS1, ALB, HNF4A, C3, ABCB4. Significantly depleted population in COVID-19 liver samples.

**BEC4:** 12.7% of Biliary Epithelial Cells. Osteopontin positive reactive cholangiocytes/HPC like cells, expressing SPP1, SOX9, LYPD6, CASR, HNF1B, ONECUT1/2, and GABRP, as well as progenitor cell response genes (ITGB6, FN14/TNFRSF12A, LTBR). Significantly depleted population in COVID-19 liver samples.

**BEC5:** 12.7% of Biliary Epithelial Cells. Immature cholangiocytes. NCAM1+ immature, reactive cholangiocytes/HPCs, expressing ITGA2, progenitor cell markers like SOX4, CK19, TROP2 and CD133. Significantly enriched population in COVID-19 liver samples.

**BEC6:** 9.6% of Biliary Epithelial Cells. Intermediate, neuroendocrine cholangiocytes. Neuroendocrine subset of cholangiocytes, expressing neural markers (TMEM132D, GRM7, HYDIN, NRXN3, LRRC4C, NTM). Trajectory analysis suggested that BEC6 may be a potential transition state between HEP1 and BEC1. Significantly enriched population in COVID-19 liver samples.

**BEC7:** 0.92% of Biliary Epithelial Cells. a minor subset of activated cholangiocytes co-expressing both epithelial and mesenchymal genes (IGFBP7, THBS2, CCBE1, COL1A2, ACTA2, EDNRA), and very active in cell cell communication, especially with the endothelial compartment (FGF, PDGF, VEGF). According to pseudotime, BEC7 is closely related to BEC6. Similar proportion between healthy and COVID-19 liver samples.

**Immune & Blood Cell Compartment**

The immune and blood cell compartment of the COVID-19 patients comprised 15.3% of all cells, spanning all major immune cell types, including macrophages/Kupffer cells (KC), T cells, B cells, Natural Killer cells and mast cells. Their cellular states may reflect the viral impact and liver homeostasis during critical illness. The immune cells were split into 16 subclusters.

**MAC1:** 22.67% of immune cells. Kupffer cells. Represents classical KC, expressing MARCO, CD163 and VISG4, as well as other major KC markers, such as MRC1, IL18, CD74, CD5L. No significant difference of proportion between COVID-19 and healthy liver samples. Significantly enriched in viral reads.

**MAC2:** 21.10% of immune cells. Intermediate, phagocytic macrophages. Lower expression of both MARCO and CD163, but increased expression of phagocytic markers including C5ARI, CPVL, RB3I and cholesterol storage genes, such as SOAT1, sphingomyelin metabolism (SPGL1) and development related cytokines (VEGF-A). Significantly enriched population in COVID-19 liver samples.

**MAC3:** 8.4% of immune cells. Proinflammatory macrophages. Characterized by an inflammatory expression program (NLRP1, HRH2, IL17RA, TNFAIP12, and VCAN), as well as markers of infiltrating monocytes like CCR2 and CD11c. No significant difference of proportion between COVID-19 and healthy liver samples.

**MAC4:** 1.79% of immune cells. Proliferating Kuppfer cells. Validated by marker expression and comparison to other murine datasets. Significantly enriched population in COVID-19 liver samples.

**NK:** 6.94% of immune cells. Natural Killer cells. Increased expression of NCAM1, CD7, NKG7, CD11a and PRF1. Significantly depleted population in COVID-19 liver samples.

**TC1:** 12.31% of immune cells. Naive CD8 T cells, with high expression of LEF1, TCF7 and GZMA, as well as CD8. Significantly depleted population in COVID-19 liver samples.

**TC2:** 3.14% of immune cells. Cytotoxic CD8+ cells with increased expression of SLC4A10, ADAM12, IL7R, CD28, CCR6 and TCF7. No significant difference of proportion between COVID-19 and healthy liver samples.

**TC3:** 2.65% of immune cells. Cytotoxic effector and memory T cells, expressing GNLY, GZMH, IFNγ, CX3CR1 and TGFBR3. Significantly enriched population in COVID-19 liver samples.

**TC4:** 0.34% of immune cells. Naive T cells with Apoptotic features, with NLRP7 and CASP8 expression, as well as high LEF1 and GZMA expression. Significantly enriched population in COVID-19 liver samples.

**BC:** 1.00% of immune cells. Mature B cells. Expressing CD20 and FCER2, CD21 and CD22. Significantly depleted population in COVID-19 liver samples.

**PC1:** 3.21% of immune cells. Terminal plasma cells, with high CD27 expression. High expression of immunoglobulins (IgJ, IKC, JCHAIN), CD38, XBP1, and ITGA8, . No significant difference of proportion between COVID-19 and healthy liver samples. Significantly enriched population in viral reads.

**PC2:** 1.00% of immune cells. Plasmablasts expressing CD38, XBP1, as well as CD70 and MYBL2, and exhibiting features of proliferation, as indicated by high MKI67 expression. No significant difference of proportion between COVID-19 and healthy liver samples.

**MAST:** 0.29% of immune cells. Mast cells, expressing both TPSB2 and CPA3. Also enriched in some neuroendocrine related markers, such as NRXN1/3, NTM and KIRREL3. Significantly depleted population in COVID-19 liver samples.

**ERY-P:** 2.90% of immune cells. Erythrocyte precursor cells, expressing a combination of hemoglobin and glycophorin genes, as well as proliferation related genes and others not present in mature red blood cells, such as CD71/ TFRC, usually only encountered in the bone marrow in adult humans. Cells of this population may be responsible for extramedullary hematopoiesis in the setting of hypoxia, as well as the modulation of the immune response during virus infection and the hematogenesis in fetal liver. Significantly enriched population in COVID-19 liver samples.

**DBL1:** 10.8% of immune cells. Doublet cluster, hepatocyte-like.

**DBL2:** 1.57% of immune cells. Doublet cluster, mesenchymal-like.

**Endothelial Cells**

The endothelial cell components in COVID-19 were substantially remodeled, accounting for 11.5% of all nuclei and spanning 12 subsets, including liver sinusoidal endothelial cells and other endothelial cell populations in an 8:1 ratio, with heavily disrupted zonation.

**EC1:** 25.2% of endothelial cells. Profibrotic LSEC-derived niche, possibly in response to systemic illness either directly or indirectly from COVID-19. These nuclei were expressing VEGFR1, FGFR1 and AKAP12, but VEGFR2 negative. Significantly enriched in COVID-19 liver samples.

**EC2:** 24.2% of endothelial cells. Typical LSEC with high LYVE1 expression, along with VEGFR2, scavenger receptors STAB1 and STAB2, MRC1, Fc-gamma receptor IIb2 (FcγRIIb2/FCGR2B), and C-type lectins (CLEC4M, CLEC4G). Significantly depleted population in COVID-19 liver samples.

**EC3:** 16.8% of endothelial cells. Profibrotic LSECs. Expression of LYVE1, and STAB2 was less high than EC2. Trajectory analysis indicates that this is a transitional state between EC2 and EC1. Significantly enriched population in COVID-19 liver samples.

**EC4:** 12.03% of endothelial cells. Classical portal endothelial cells in hepatic arteries and veins. Low expression of LYVE1, STAB2 and CD32B. Expression of both EFNB2 and EDNRB, suggesting periportal localization and association with NO production and vasodilation. Similar proportion between COVID-19 and healthy liver samples.

**EC5:** 8.56% of endothelial cells. PLVAP Fibroic LSECs. Classical endothelial cells in hepatic arteries and veins. High level of PECAM1, vWF, CD34, CD39, PDGFD. Significantly depleted population in COVID-19 liver samples.

**EC6:** 4.08% of endothelial cells. Central Venous endothelial cells. High expression of PECAM1, EGFL7, but also numerous MT genes, as well as SERPINA1 and HBA2. Similar proportion between COVID-19 and healthy liver samples.

**EC7:** 3.53% of endothelial cells. EMT related endothelial cells. Convolved transcriptional profile of both endothelial and biliary epithelial markers (CFTR, THBS2, CCBE2, DGKB). While this transition has been shown in cancer, this is potentially a doublet cluster. Significantly enriched in viral reads. Significantly enriched population in COVID-19 liver samples.

**EC8:** 1.79% of endothelial cells. Features of classical vascular endothelial cells, expressing ENTPD1, vWF, RSPO3, but also EDN1, NOS1 and COX1, and the anti-inflammatory gene C7. Significantly depleted population in COVID-19 liver samples.

**EC9:** 1.25% of endothelial cells. Potentially a subpopulation of infiltrative monocyte cells, or pro-inflammatory endothelial cells with medium LYVE1 expression and very high ICAM1 expression, but also expression of PTPRC, CD44, IFNGR2, FCGR3A and ADA2, relating to the IL-2 pathway. Significantly enriched population in COVID-19 liver samples.

**EC11:** 0.79% of endothelial cells. Rare subset of VEGFR1 negative cells, expressing VEGFR3, representing lymphatic endothelial cells. Also expressed lymphatic markers like PDPN and PROX1. Detected almost exclusively in COVID-19 liver samples, but no significant difference.

**EC12:** 0.70% of endothelial cells. Possibly a subpopulation of replicating endothelial cells, observed in other murine studies, with high expression for both proliferation (MKI67, TOP2A, CENPF, ANLN) and angiogenesis-associated genes. Detected almost exclusively in COVID-19 liver samples, but no significant difference.

**DBL3:** 0.98% of endothelial cells. Hepatocyte features**.** Potentially technical artifact.

**Mesenchymal Cells**

The mesenchymal cell component of the COVID-19 nuclei accounted for 5.8% of all the nuclei and represented all major cell lineages: quiescent and activated hepatic stellate cells (HSC), smooth muscle cells, myofibroblasts and fibrocytes and was largely affected by COVID-19, consistent with a profibrotic activation phenotype.

**MES1:** 26.31% of mesenchymal cells. Represents quiescent HSCs (qHSCs), distinguished by pericyte lineage markers (RELN. HGF and COL25A1). Trajectory analysis strongly supported a MES1, MES2, MES3, MES8 transition in continuous time, consistent with lobular to portal migration. No significant difference between proportions in COVID-19 and healthy liver samples.

**MES2:** 22.9% of mesenchymal cells. Partially activated HSC (aHSC), with distinct markers like CBE1, RXFP, and PTGIS, as well as genes related to collagen receptors and chemokine signaling being highly expressed. Significantly enriched population in COVID-19 liver samples.

**MES3:** 22.4% of mesenchymal cells. ECM-associated HSC subset, expressing both genes involved in collagen receptor signaling pathways over production (ITGA2, ITGA11, C7, CCBE1, NFASC, EPHA3, COL1A2, CACNA1C, and COL4A4). Mapped to the portal tract. Significantly enriched population in COVID-19 liver samples.

**MES4:** 8.04% of mesenchymal cells. Smooth Muscle Cells (SMCs), with a typical transcription program comprising of EBF1, SLIT3, PLA2G5, PDE3A , ID4 and TRPC4, as well as *bona fide* SMC differentiation drivers MYH11 and MYOCD. Significantly enriched population in COVID-19 liver samples.

**MES5:** 7.05% of mesenchymal cells. Inactive, bone marrow derived fibrocytes, based on the co-expression of monocyte markers CD45, CD11b and ACTA2, as well as collagen gene expression and high GALNT17. Overally, it had lower expression of mesenchymal markers compared to HSC subpopulations (MES1-3). Mapped to the portal tract. Significantly depleted population in COVID-19 liver samples.

**MES7:** 5.91% of mesenchymal cells. High expression of MT genes and low nuclear mRNA counts, indicating that it might be an artifact. Significantly depleted population in COVID-19 liver samples.

**MES8:** 0.65% of mesenchymal cells. Minor subset of activated, periportal myofibroblasts, characterized by high expression of EMILIN1, BGN, COL3A1, TAGLN, HGF, COL1A1, COL1A2, COL4A4, COL6A1, RWRN1, CXCL12, TIMP1, CTGF, DCN and C7. Was very active in CCI analysis, with many interactions between MES8 and HEP7. Mapped to the portal tract. Significantly depleted population in COVID-19 liver samples.

**DBL4:** 6.71% of mesenchymal cells. Doublets with hepatocyte features.
