## Extended Data Table 1 for "A single-nucleus and spatial transcriptomic atlas of the COVID-19 liver reveals topological, functional, and regenerative organ disruption in patients"

**Extended Data Table 1:** Liver serum markers for COVID-19 and control liver samples.

| **Donor** | **ALT (IU/L)** | **AST (IU/L)** | **ALP (IU/L)** | **T. Bilirubin (mg/dL)** |
| --- | --- | --- | --- | --- |
| L1 | 11 | 35 | 66 | 0.4 |
| L2 | 63 | 92 | 69 | 0.7 |
| L3 | 70 | 378 | 156 | 0.4 |
| L4 | 59 | 183 | 172 | 0.5 |
| L5 | 173 | 105 | 185 | 3.5 |
| L12 | 109 | 389 | 118 | 4.8 |
| L13 | 64 | 130 | 127 | 1.1 |
| **Controls** | **ALT (IU/L)** | **AST (IU/L)** | **ALP (IU/L)** | **T. Bilirubin (mg/dL)** |
| L18 (C41) | 182 | 69 | 76 | 4 |
| L19 (C58) | 29 | 44 | 73 | 13 |
| L20 (C70) | 41 | 26 | 89 | 4 |
| L21 (C72) | 57 | 48 | 55 | 5 |
| **Normal range** | 10 - 50 | 10 - 50 | 35 - 130 | 0.0 - 1.0 |
