## Extended Data Table 6 for "A single-nucleus and spatial transcriptomic atlas of the COVID-19 liver reveals topological, functional, and regenerative organ disruption in patients"

**Extended Data Table 6:** Summary of viral loads using RT-PCR in donors L1-5 who expired due to COVID-19 (LOQ: limit of quantification).

|  | **SARS-CoV-02 Viral Load**  **(copies/μg total RNA)** | **Log10 viral loads genomic** | **Log10 viral load subgenomic** |
| --- | --- | --- | --- |
| L1 | 7,783 | 3.89 | 3.16 |
| L2 | 2 | 0.30 | < LOQ |
| L3 | 0 | < LOQ | < LOQ |
| L4 | 2 | 0.30 | < LOQ |
| L5 | 1 | < LOQ | < LOQ |
