## Extended Data Table 8 for "A single-nucleus and spatial transcriptomic atlas of the COVID-19 liver reveals topological, functional, and regenerative organ disruption in patients"

**Extended Data Table 8:** Summary of liver histopathology findings for samples L1 to L4. H&E staining, CK19, and α-SMA IHC, as well as connective tissue staining (picrosirius red) were performed in four consecutive core biopsies samples and evaluated by an expert clinical liver pathologist (I.N.).

|  | **Steatosis** | **Inflammation** | **Activation of stellate cells** | **Ductular reaction** | **Fibrosis** | **Other notes** |
| --- | --- | --- | --- | --- | --- | --- |
| L1 | Moderate microvesicular steatosis | Focal mild portal mononuclear inflammation including plasma cells | Focal zone 3 myofibroblastic proliferation | Minimal focal bile duct proliferation | Centrivenular and focal periportal fibrosis (stage 0-1) | Moderate hemosiderin, Kupffer cell deposition |
| L2 | Mild macrovesicular steatosis | Focal mild portal mononuclear inflammation including plasma cells | Focal zone 1 and 3 myofibroblastic proliferation | Areas of bile duct proliferation | Centrivenular, sinusoidal and focal periportal fibrosis (stage 2) | Focal hepatocellular centrivenular necrosis |
| L3 | Minimal macrovesicular steatosis | Focal mild portal mononuclear inflammation | Diffuse panlobular myofibroblastic proliferation | Extensive diffuse bile duct proliferation | Centrivenular, sinusoidal and periportal fibrosis with bridging and focal nodule formation (stage 3-4) | none |
| L4 | none | Focal mild portal mononuclear inflammation | Prominent predominantly zone 3 myofibroblastic proliferation | Prominent, multifocal bile duct proliferation | Centrivenular, sinusoidal and periportal fibrosis and focal nodule formation (stage 3-4) | Hemosiderin-laden Kupffer cells |
